# Honey Bee Diversity is Swayed by Migratory Beekeeping and Trade Despite Conservation Practices: Genetic Evidences for the Impact of Anthropogenic Factors on Population Structure

**DOI:** 10.1101/154195

**Authors:** Mert Kükrer, Meral Kence, Aykut Kence

## Abstract

Intense admixture of honey bee (*Apis mellifera* L.) populations is mostly attributed to migratory beekeeping practices and replacement of queens and colonies with non-native races or hybrids of different subspecies. These two practices are also heavily carried out in Anatolia and Thrace where 5 subspecies reside naturally.

Here, we carried out an analysis of population structure of honey bees sampled from six different regions (n = 250) in order to test the genetic impacts of migratory beekeeping, queen and colony trade and conservation efficacy of isolated regions. A total of 30 microsatellite markers were used in four multiplex reactions.

Direct genetic impact of migratory beekeeping was demonstrated first time based on a comparison of assignment of individuals to their geographically native populations where migratory colonies showed less fidelity. We found genetic evidence for them acting as a hybrid zone mobile in space and time, becoming vectors of otherwise local gene combinations.

The effects of honey bee trade were revealed by the presence of very high introgression levels from the highly commercial Caucasian bees naturally limited to a narrow range. We also measured the direction and magnitude of this gene flow connected with bee trade.

Comparison between regions that are either open to migratory beekeeping or not let us evaluate the status of isolated regions as centers of limited gene flow and showed the importance of establishing such regions.

Despite signs of gene flow, our findings confirm high levels of geographically structured genetic diversity of four subspecies of honey bees in Turkey and emphasize the need to develop policies to maintain this diversity.

Our overall results might potentially bear a wider interest to the community since they constitute an important attempt to quantify the effects of anthropogenic impacts on established patterns of honey bee diversity. Our measurable and justified findings on migratory beekeeping, queen and colony replacements as well as conservation implications will hopefully be of use for the decision makers and other stakeholders.

## Introduction

The Western honey bee, *Apis mellifera* L., is a species which plays role together with other pollinators in pollination of wild and cultivated plants while the species also have significant economic importance in terms of honey and other bee products output (Morse 1991; Breeze et al. 2011). In addition to its ecological and economic importance, it is a model study organism both for evolution of eusociality and sophisticated cognitive abilities (Weinstock et al. 2006).

Natural distribution of A. mellifera includes Central and Southwest Asia, Europe and Africa but the species was also introduced to East and Southeast Asia, Australia and the Americas mainly on purpose for its economic benefits (Ruttner 1988). Morphological and molecular studies point to four major lineages of numerous –more than 20- subspecies (Ruttner, 1988; Whitfield et al. 2006). The four widely recognized lineages are A (Africa), M (western and northern Europe), O (Near East and Central Asia) and C (Eastern Europe) lineages.

Although bearing controversies, studies with Single Nucleotide Polymorphisms (SNPs) in the past decade supported the hypothesis that *A. mellifera* have originated in the tropics or subtropics in Africa and colonized its natural range by two main routes: one through Gibraltar and one through Suez and then Bosphorus, ending up with a secondary contact between the highly divergent A and C lineages around Alps (Whitfield et al. 2006; Han et al. 2012; Walberg et al. 2014; Harpur et al. 2014; Cridland et al. 2017).

Both the honey bees and wild pollinators are thought to be on decline (locally and/or globally depending on the species and region of concern) due to factors some of them relating closely to human activities. Among them, destruction and fragmentation of natural habitats, toxicity caused by pollution and pesticides –as such widely used neonicotinoids-, diseases and their spread getting easier, invasive species are leading the way (Meffe 1998; Brown & Paxton 2009; Van Engelsdorp & Meixner 2010; Blacquiere et al. 2012). Honey bees also, especially wild populations that are not managed by beekeepers (including the feral populations), take their share from the situation like the other species in the genus *Apis* –namely *Apis cerana, Apis florea, Apis dorsata* and other native bees of Asia (Oldroyd 2007; Dietemann et al. 2009; Van Engelsdorp et al. 2009; Genersch 2010; Evans & Schwarz 2011).

Besides such negative consequences created by human activities; the genetic admixture of honey bee populations due to bee trade, including complete replacement of local bees with non-natives and beekeeping practices involving movement of colonies from one region to the other impose another kind of pressure on the species: the loss and/or swamping of locally adapted gene combinations and local or global extinctions of native honey bees (De la Rua et al. 2009).

All those factors and their interactions, including genetic and environmental ones, when combined, may have an increased adverse effect on honey bees and may be the reasons behind continuous or discrete events of sudden colony losses with rapid depletion of worker bees while the queen continues laying eggs accompanied by lack of dead bees in and around the hive; the syndrome called as Colony Collapse Disorder (CCD) or Colony Depopulation Syndrome (CDS) (Van Engelsdorp et al. 2009; Neumann & Carreck 2010).

Resilience of the honey bees may be lying in the adaptations they accumulated over thousands of years, and new potentials reside in their genetic diversity. It is highly probable that a combination of many above mentioned factors/threats are taking their places in the recent declines by weakening the colonies step by step. Due to altered rankings of performance of subspecies in varying environments, it is generally accepted that honey bees’ resistance or tolerance to these factors differ greatly and locally adapted variants may be encountering less stress, thus remain standing strong (Büchler et al. 2015). Hence, research on honey bee diversity in the global context and at various levels (genetic, individuals, colonies, populations, ecotypes and subspecies) is of great importance for maintaining the species’ and ecosystem services they provide as well as their economic usefulness.

In recent years’ research conducted on honey bee population structure in European countries, it was shown that the past structure was lost or strongly disturbed (Dall’Olio et al. 2007; Canovas et al. 2011; Bouga et al. 2011). Introgression of non-native DNA was monitored in wild populations of Sudan (El-Niweiri & Moritz 2010). Among the anthropogenic effects, mainly queen and colony trade and replacement of native honey bees with non-natives as well as migratory beekeeping were the usual suspects.

Despite grounded suspicions there are very few studies that investigate and test the direct genetic consequences of human practices on honey bee diversity. Therefore, the aims of this series of experiments were testing different hypotheses about recent heavy/any admixture of honey bee populations across four subspecies by making use of microsatellite markers as well as i) evaluating the status of isolated regions as a conservation implication where migratory beekeeping is prohibited, restricted or very scarce due to lack of preference of migratory beekeepers or attitude of local beekeepers ii) acquiring and demonstrating the direct genetic outcomes of migratory beekeeping by a series of comparisons between migratory and stationary colonies iii) seeking for the effects of unregulated queen and colony trade by figuring out the origin, extent and direction of introgression between populations.

With five subspecies dwelling within its borders and with a variety of beekeeping strategies, Turkey makes a good stage for chasing genetic evidences for the impact of anthropogenic factors on one of the most important crop and wild plant pollinators. Beekeeping is an old tradition in those lands which dates back to 6600 BC and Hittites civilization (Akkaya & Serhat 2007), while still intensively practiced in Turkey where there are more than 8 million hives distributed all over the country. This is the third highest number in the world, alone tripling those of the USA and reaching the half of the EU total (USDA NASS 2019, European Parliament 2017).

Corresponding to one-fourth to one-fifth of all recognized subspecies of *A. mellifera; A. m. meda, A. m. syriaca, A. m. caucasica, A. m. anatoliaca* from the O-lineage and an ecotype from C subspecies group exist in Turkey (Kandemir et al. 2005). Even A-lineage genetic material was also characterized in native bees from the Levantine coast of Turkey (Kandemir et al. 2006) bringing together genetic elements from three continents. Major subspecies found in and around Anatolia are shown in Fig. 1a.

**Figure 1.**
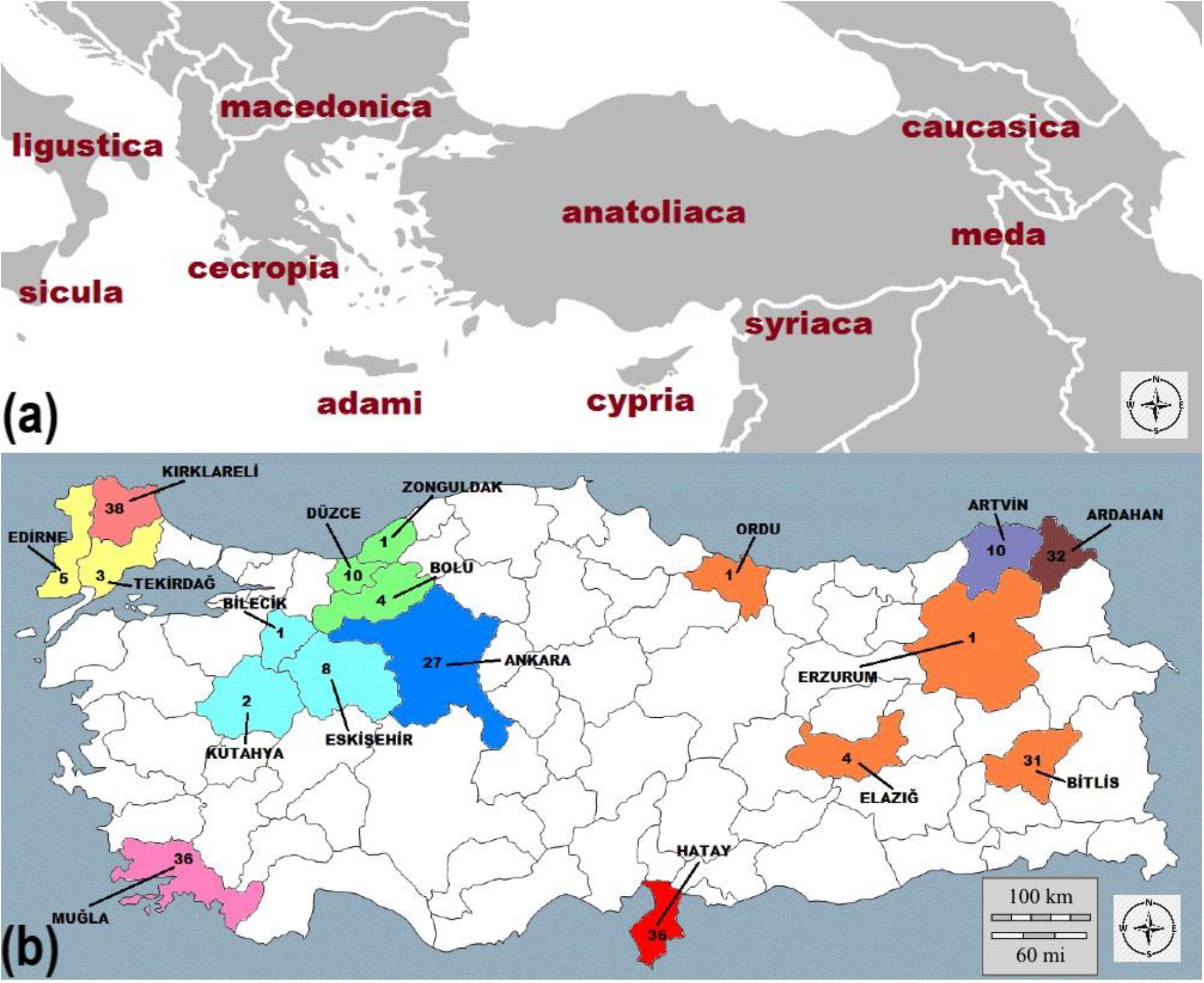
Geographic distribution of (a) major honey bee (*A. mellifera*) subspecies in and around Anatolia (b) sampling sites and sample sizes.

Anatolia and Thrace, when taken together, harbor a vast diversity: honey bees belonging to three different lineages meet, exchange genes and adapt to local conditions determined by diverse climatic, topographical and floristic variations available (Bouga et al. 2011). Refugial status of Anatolia during the ice ages contributed present enhanced levels of biodiversity (Hewitt 1999). Studies concerning honey bee populations of Turkey (Bodur et al. 2007; Kence et al. 2009) demonstrated high genetic structuring among them and confirmed the presence of divergent populations pointing to different subspecies. They, all together, drew attention to this rich diversity hotspot present and particularly under threat in Anatolia and Thrace as well as importance to its conservation.

Despite that, still prevail the arguments in popular opinion -with significant effect on decision makers and other stakeholders- that honey bee ecotypes are inevitably lost due to gene flow facilitated by anthropogenic factors, so the relevance of investing in a strategy involving conservation of locally adapted variants are unremittingly questioned. This study aimed to quantify and weigh the impacts of anthropogenic factors and conservation efforts on the present condition of honey bee genetic diversity.

## Materials and Methods

### Sampling

We sampled a total of 250 honey bees each from different colonies from 18 provinces during the period of March 2010 and August 2012. Of those 250 honey bees, 174 were from apiaries that were stationary and 76 were from migratory ones. Beekeepers declared that they used original honey bees from stocks native to the area and that they did not purchase non-native queens or colonies in the last ten years.

We grouped samples from provinces with small sample sizes together with nearby provinces to form 10 major localities: Kırklareli, Edirne+ (Edirne and Tekirdağ), Muğla, Eskişehir+ (Eskişehir, Kütahya and Bilecik), Düzce+ (Düzce, Zonguldak and Bolu), Ankara, Hatay, Bitlis+ (Bitlis, Elaziğ, Erzurum and Ordu), Ardahan, and Artvin. Those localities correspond to natural distribution range of four subspecies. Those subspecies are *A. m. syriaca* in Hatay, *A. m. caucasica* in Ardahan and Artvin, *A. m. anatoliaca* in Düzce, Eskişehir+, Muğla and Ankara from the O lineage and an ecotype from C subspecies group in Kırklareli and Edirne+ by excluding the fifth subspecies *A. m. meda*. We carried out combinations of locations according to geographical proximity; similarity in terms of climatic, topographic and floral variables; results of previous studies as well as results of preliminary analysis of this study. Sampling sites and sample sizes can be seen in Fig. 1b.

The samples were kept in −80 °C until genetic analysis.

### Genotyping

We isolated DNA from bee heads by QIAGEN DNeasy Blood and Tissue Kit following the procedure of the producer for insect samples with slight modifications. We grouped a set of 30 microsatellite loci into four clusters for two 7-plex (<u>set 1:</u> AP218, A113, AB024, AP249, A088, AP001, AP043 and <u>set 2:</u> AP049, AP238, AC006, AP243, AP288, HBC1602, A107) and two 8-plex (<u>set 3:</u> A079, AC306, AP226, A007, HBC1601, AP068, A014, AP223 and <u>set 4:</u> AP019, AB124, A043, A076, AP273, AP289, HBC1605, A028) polymerase chain reactions (Estoup et al. 1995; Solignac et al. 2003; Bodur et al. 2007; Shaibi et al. 2008; Tunca et al. 2009). A software program, Multiplex Manager 1.2 (Holleley & Geerts 2009), was used for constructing the multiplex groups. Information on primer pairs, fluorescent dyes and PCR conditions are provided in the supplementary file.

Detection of microsatellite allele sizes was achieved by capillary electrophoresis with ABI 3730XL sequencing machines. We were not able to amplify locus A076 consistently across the samples thus we definitely excluded it from the data set and the downstream analysis.

### Population structure

We calculated pairwise F_ST_ values by Arlequin 3.5 (Excoffier et al. 2005), Mantel test was applied to account for isolation by distance procedure. Pairwise population distances were calculated (Reynolds et al. 1983) by Populations 1.2.32 software (Langella 2011) and visualized by the online tool Interactive Tree of Life v4 (Letunic and Bork 2019). We used PAST4 and PCAgen software to plot populations on a two-dimensional space by a PCA based on correlation matrix between groups (Goudet 1999; Hammer 2001).

Population structure was estimated by Structure 2.3.3 (Pritchard et al. 2000), K values of distinct populations were analyzed by Structure Harvester software (Earl & von Holdt 2012), and we used Clumpp software (Jakobsson & Rosenberg 2007) to permute the membership coefficients of individuals determined by Structure 2.3.3 and Distruct software to (Rosenberg 2004) visualize the results obtained by Clumpp.

Other population genetic parameters and diversity indicators were also estimated and they are provided as supplementary file. These parameters and indicators contain frequency of null alleles, allelic richness and diversities, inbreeding and prevalence of close relatives, number effective alleles, levels of heterozygosity, deviations from Hardy-Weinberg and linkage disequilibrium, bottlenecks, effective population sizes and microsatellite information index.

### Statistical analyses

We then used membership coefficients obtained, to test hypotheses about beekeeping practices, isolated regions and queen/colony trade. For the analysis, we first arcsine root square transformed the coefficients since the data was composed of proportions and non-normally distributed. Then we carried out Shapiro, Mann-Whitney *U*, Kruskal-Wallis, Dunn’s, *F*, ANOVA, Tukey’s and *t* tests wherever necessary and applicable to compare mean membership coefficients and estimated Cohen’s *d* to determine effect sizes. Those were carried out in R statistical software using packages pwr, effsize, dunn.test and dabestr (R Core Team 2013, Torchiano 2016; Dinno 2017; Champely et al. 2018; Ho et al. 2019). R code is provided as a supplementary material.

We made use of estimation plots to visualize untransformed data for membership coefficients and impact of various factors on them. This is a less conventional method when compared to bar or boxplots and reporting of significance tests but much more convenient and powerful method to summarize the whole data in an unbiased way by displaying all measurements and effect sizes as well as precision of estimates and distribution of mean differences (Ho et al. 2019).

### Beekeeping practice: migratory vs stationary

For the first hypothesis to be tested, we compared membership coefficients of migratory and stationary colonies in Ankara, Muğla and Hatay separately, for the three provinces combined and for the total data set. If the migratory colonies acted as a potential vector of foreign alleles then they would have much lower probabilities of being assigned to their own clusters.

### Isolated regions as a conservation practice

The second hypothesis was about isolated regions. If the isolated regions were efficient in preserving genetic diversity by preventing gene flow between different clusters then one would expect to see higher membership coefficients for stationary individuals belonging to these regions and lower for stationary individuals that belong to regions open to migratory beekeeping.

Kırklareli is a province that is declared officially as an isolated region where migratory beekeepers could not visit for years at first thanks to local beekeepers’ negative attitude towards them. The region is home to a Carniolan ecotype carefully maintained by local beekeepers. Ardahan is legally declared a conservation and breeding area for *A. m. caucasica* so migratory beekeepers cannot enter the province and queen import from other subspecies is forbidden. Parts of Artvin province are also officially declared as isolated regions for conservation of *A. m. caucasica* as a pure race. The province in general is rarely visited by migratory beekeepers for geographical reasons and beekeepers there, dealing with mass queen breeding, do not use non-native queens. We compared these three provinces with the other six regions (Edirne+, Muğla, Düzce+, Eskişehir+, Ankara and Hatay) where migratory beekeeping and bee trade are freely exercised.

### Effect of queen and colony trade

Third set of tests were about the impacts of honey bee trade. We compared the estimated proportion of genomes assigned to a different cluster than the native cluster among individuals of the total data set to find out which cluster contributed most to other clusters’ gene pools.

Ardahan and Artvin provinces host the *A. m. caucasica* subspecies which is also widely used for commercial purposes and the *caucasica* queens and their hybrids are sold all over the country. But these provinces are also limited to a very narrow range in the Northeast of the country and are declared isolated regions. So, a possible high introgression of their alleles would mostly, if not completely, be due to replacement of queens and colonies.

We also investigated further patterns across populations to understand the magnitude and direction of the gene flow by tracing the signs of those misassigned proportions within localities.

## Results

We calculated FST values by using both the frequencies obtained in the study and by using the null allele corrected frequencies. We calculated for the stationary (n = 174) colonies an overall FST of 0.065 and an F_ST_ of 0.067 after null allele corrections. For migratory colonies the values were 0.011 and 0.015 respectively and for all the 250 samples the values were 0.046 and 0.047.

Phylogenetic tree we constructed by using pairwise population distances based on stationary colonies only resolves four distinct branches corresponding to four subspecies (Fig. 2b). Thracian samples constitute the extreme end of the unrooted tree. The other end is divided to three almost equidistant branches of Caucasian, Levantine and Anatolian samples.

**Figure 2.**
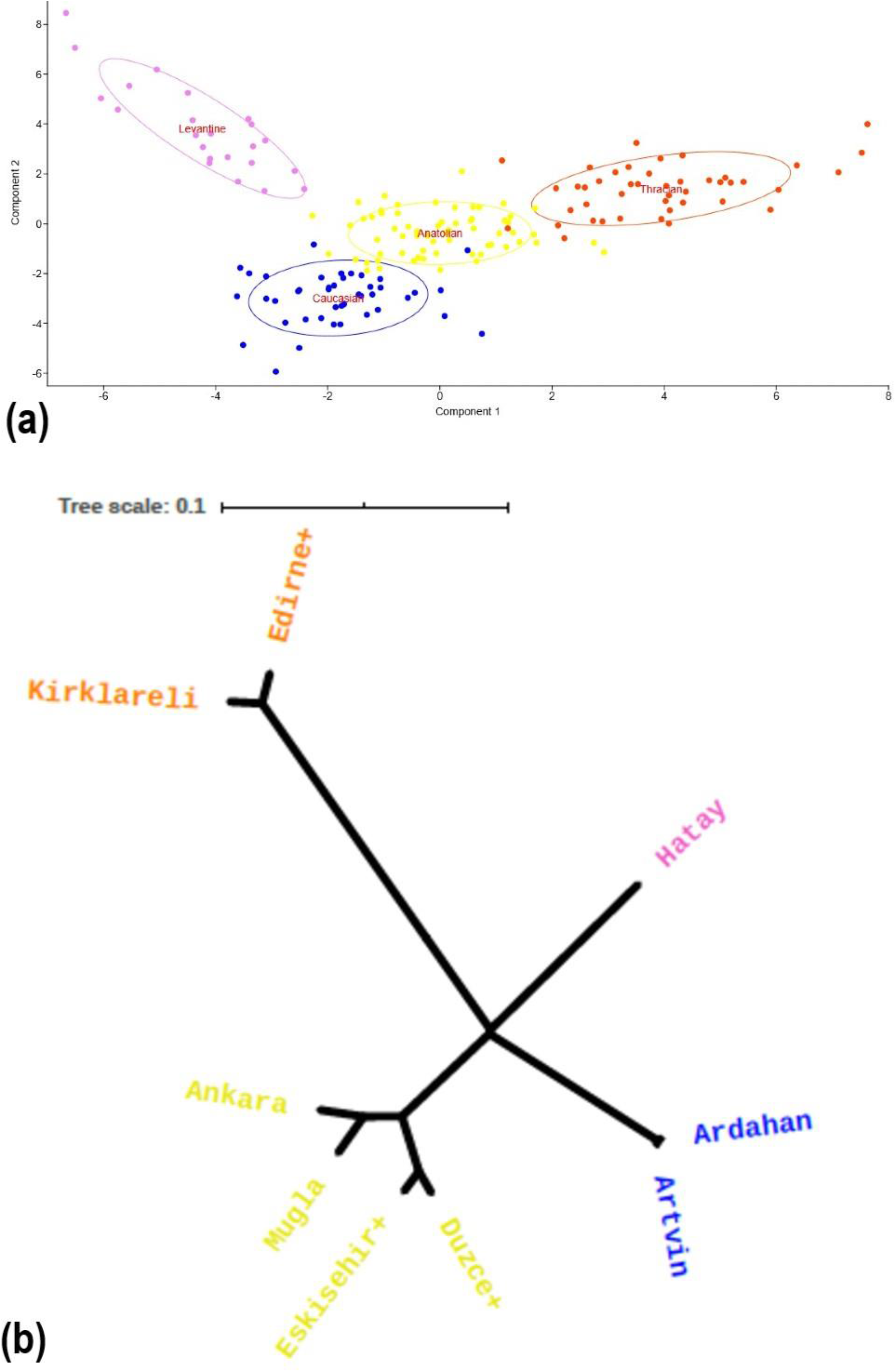
(a) PCA of stationary colonies, Component 1: 41.8%, Component 2: 32.1% (b) UPGMA tree of honey bee populations based on Reynolds’ 1983 genetic distances (orange: Thracian, yellow: Anatolian, blue: Caucasian, violet: Levantine clusters).

We plotted stationary colonies, migratory colonies and the overall data for the regions of sampling on 2D spaces by carrying out Principle Component Analysis (Fig. 2a) which showed a similar pattern with the UPGMA tree. First axis designating the first principle component differentiated samples those in Thrace whereas the second one corresponding to the second component differentiated subspecies in Anatolia (*syriaca, anatoliaca* and *caucasica*). The x and y axes explained 41.8% and 32.1% of the variance within the samples.

Genetic distances in stationary colonies showed significant correlation (p < 0.001) with geographic distance but those of migratory colonies were not correlated with geographic distances. Results of Mantel test point to an isolation by distance pattern in stationary colonies that is lost in migratory ones.

Concerning the Structure results, the best K values were selected by the Structure Harvester program as 2 and 4 with similar outcomes, K=2 being slightly likelier than K=4 which hint for lineage level diversification of C and O ancestries. We calculated membership coefficients of individuals to the observed clusters in K=4 since it can be biologically attributed to relevant subspecies under investigation and we used them for further hypothesis testing. Clustering analyses showed no population structuring for migratory colonies (Fig. 3a) in contrast to stationary colonies and the overall data (Fig. 3b and 3c). Concerning the overall data, however, distortion in the population structure caused by migratory colonies is evident in higher admixture levels observed.

**Figure 3.**
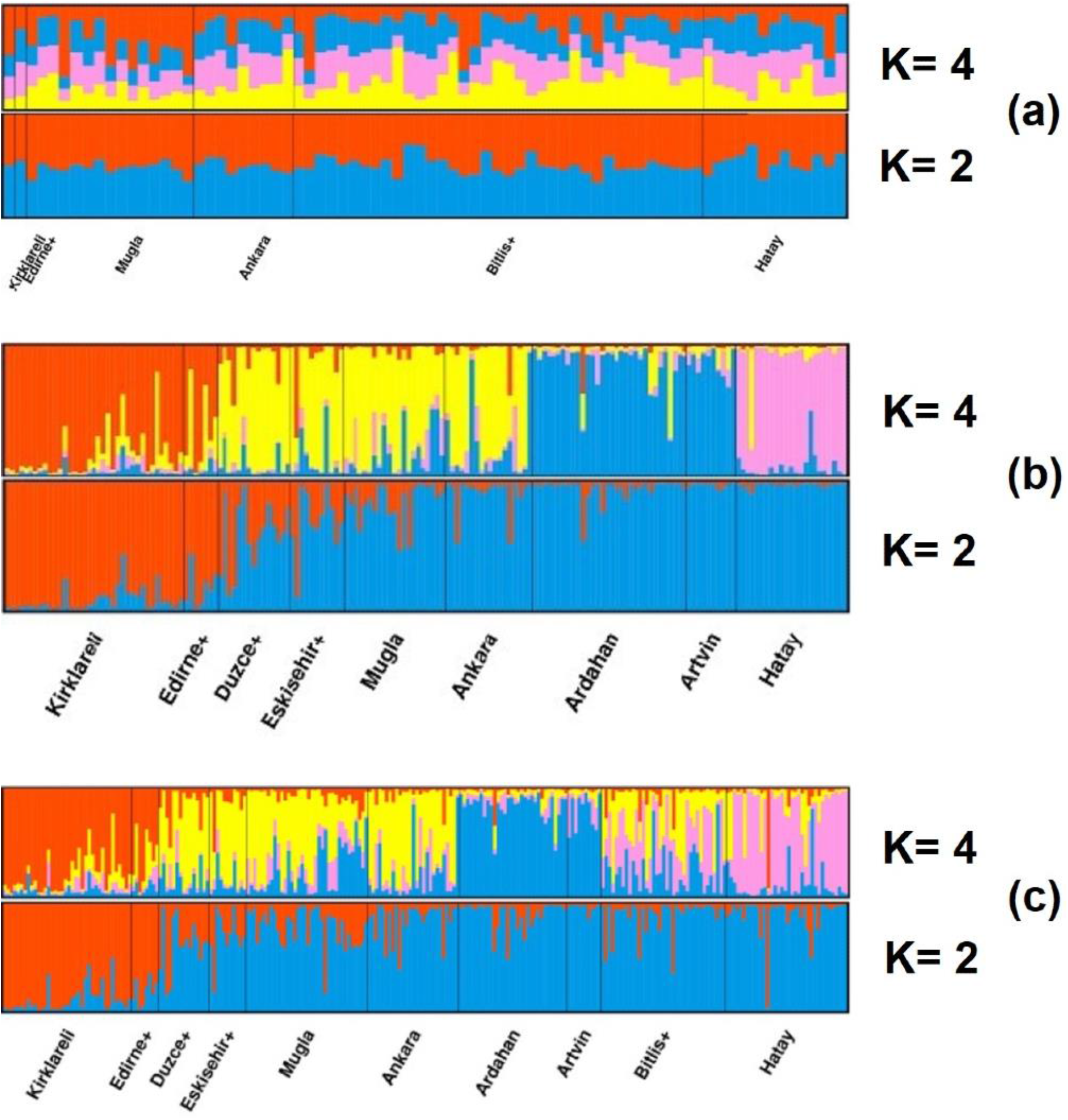
Estimated population structure and clustering of honeybees in Anatolia and Thrace for (a) migratory colonies (b) stationary colonies (c) the whole sample (orange: Thracian, yellow: Anatolian, blue: Caucasian, violet: Levantine clusters).

We compared individuals from stationary and migratory colonies according to their membership coefficients belonging to their native clusters (or it can be called their expected natural populations alike). The mean values and effect sizes as well as the significance level of the differences were summarized in Table 1. Boxplots contrasting the arcsine root square transformed membership coefficients for migratory and stationary colonies are shown in Fig. 4a and scatter plots are very much similar but visualizing raw membership coefficients for each sample are shown in Fig. 4b.

**Table 1.**
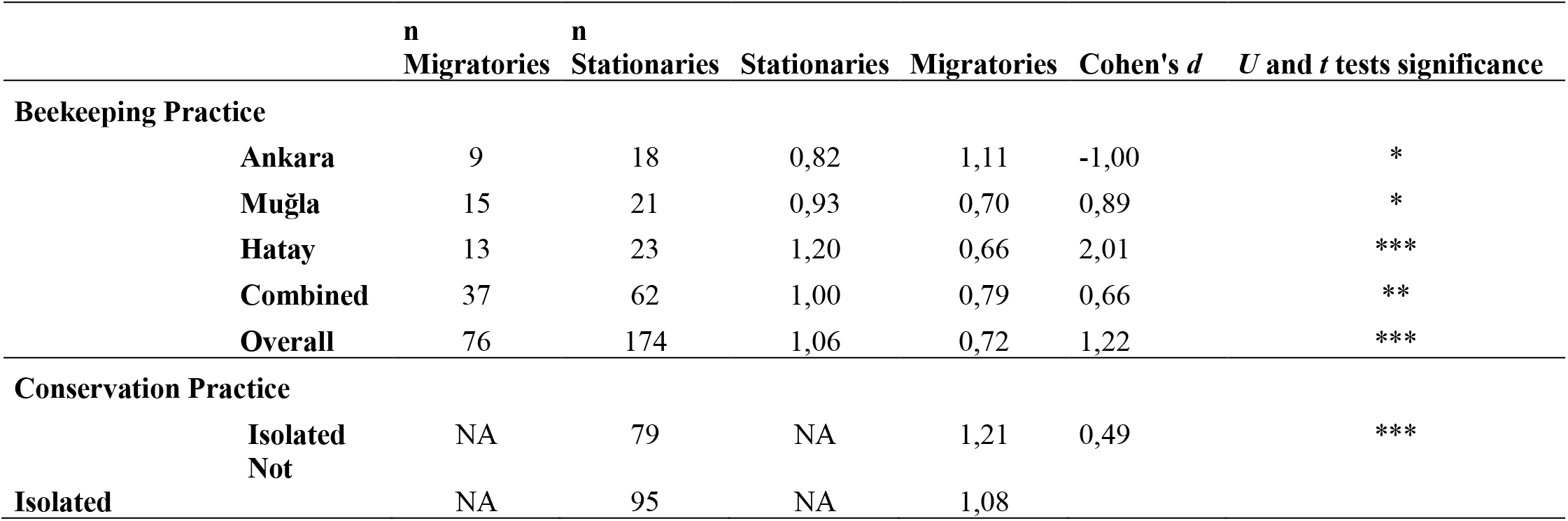
Genetic impact of beekeeping and conservation practices on (arcsine root square transformed) membership coefficients to native clusters (** *p*<0.05, \*\**p*<0.01 and \*\*\**p*<0.001).

**Figure 4.**
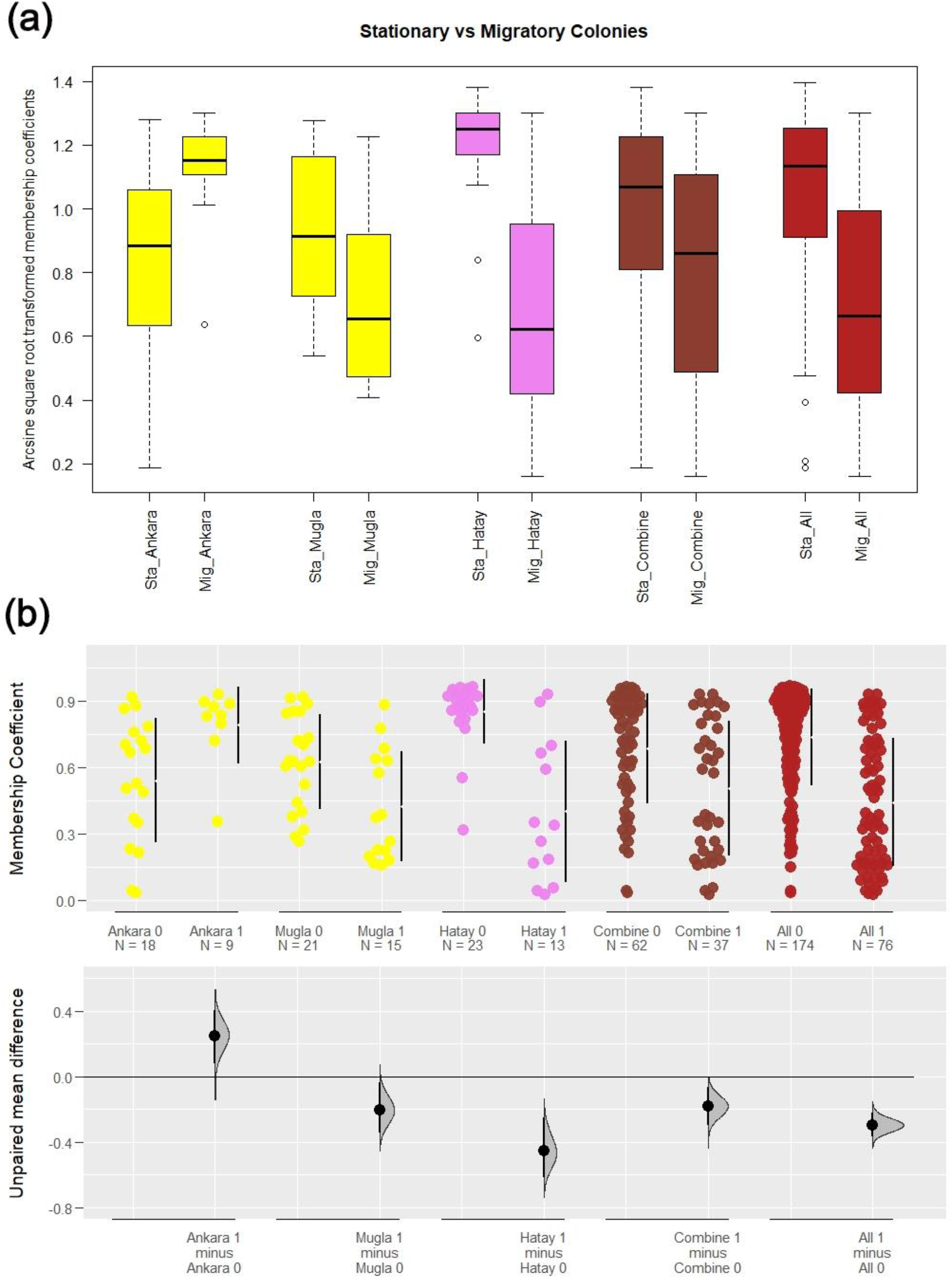
Comparison between stationary (Sta_) and migratory (Mig_) colonies in Ankara, Muğla and Hatay, as well as these three provinces combined and the whole data set, n = 250 (a) boxplot display of arcsine root square transformed membership coefficients (b) scatter plot with estimations of mean differences based on raw individual membership coefficients (yellow: Ankara and Muğla belonging to Anatolian cluster, violet: Levantine cluster, coral: for a combination of three provinces, firebrick: whole data).

Estimation plots not only fairly visualize the real distribution of the data but also let us compare the effect sizes and their precision. Stationary colonies are annotated as <Group name> 0 and migratory colonies are as <Group name> 1 (Fig. 4). Bars right to the data points refer to the 25% and 75% quartiles and the gap between them is the median value for the sample. The zero line below correspond to the mean membership coefficients of stationary colonies in each pairwise comparison. The Euclidean distances from those means for the migratory colonies are shown as dots with a 95% confidence interval bar around. Also, distributions of the estimation statistics are included. So that we can comprehensively compare the strength of the drift for different populations and subsets of the data.

Stationary colonies from Muğla and Hatay were quite more likely to be assigned to their own clusters than the migratory colonies from these provinces, the same held when we compared the combined data from the three provinces or all the migratory and stationary colonies. However, the situation was the reverse in Ankara possibly due to factors we discuss below. Stationary colonies from that province reflected patterns of high admixture. The difference between stationary colonies and migratory in Muğla are much less when compared against the ones in Hatay, signaling for a possible higher level of admixture in Muğla.

For that first comparison we used the complete (n = 250) data set to be able to quantify the differences in membership coefficients for migratory and stationary colonies. But for the rest of the analysis we used the subset of data which is only composed of stationary colonies (n = 174) since this would better reflect the population structure.

In the first scatter and the corresponding boxplots (respectively Fig. 5b for raw membership coefficients and Fig. 5a for transformed values) one can observe that within each locality samples are assigned with high proportions to their native clusters despite some admixed individuals. Also, one can see through observation of unpaired mean differences that Kırklareli, Ardahan, Artvin, Hatay and to a lesser extent Düzce play role as centers of genuine subspecies diversity with exceptionally high levels and few individuals of admixed origin.

**Figure 5.**
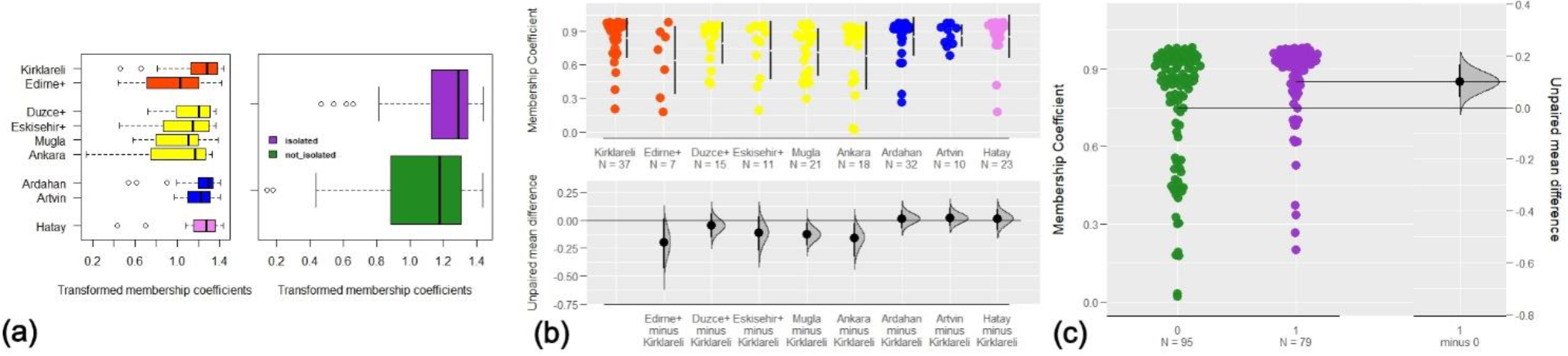
Comparison between isolated regions and regions that are open to migratory beekeeping (a) first boxplot display the arcsine root square transformed membership coefficients for 9 populations whereas the second one presents a comparison of samples within isolated regions and those are not (b) scatter plot with estimations of mean differences based on raw individual membership coefficients to the native clusters (c) scatter plot contrasting individual raw membership coefficients with an estimation of mean differences (orange: Thracian, yellow: Anatolian, blue: Caucasian, violet: Levantine clusters, orchid and “1”: isolated regions, green and “0”: regions open to migratory beekeeping).

But when we compared isolated regions (Kırklareli, Ardahan, Artvin) and regions open to migratory beekeeping (consisting of Edirne+, Muğla, Ankara, Düzce+, Eskişehir+ and Hatay provinces in our sample) in terms of their arcsine transformed membership coefficients (Table 1 for means, effect sizes and significance of the difference and Fig. 5a second boxplot) we witnessed that -as expected-stationary colonies within isolated regions showed significantly higher fidelity to the original clusters. This is also obvious in the estimation plot in Fig. 5c where the mean membership coefficients of samples that are from regions open to migratory beekeeping (green colored group designated as <0>) fall beyond the 95% confidence interval of the estimated mean of the samples from the isolated regions (orchid colored group designated as <1>).

Even if the individuals are assigned with high probability to their own clusters, let’s say with a 90% of probability, this means that 10% of their genome still belongs to other clusters. Given that there are four clusters, we investigated if any of these misassigned genome parts were enriched for any of them. Mean transformed values for Thracian cluster misassignments among individuals of the other populations were 0.16 and 0.25 for Anatolian cluster, 0.26 for the Caucasian and 0.20 for the Levantine (Fig. 6a).

**Figure 6.**
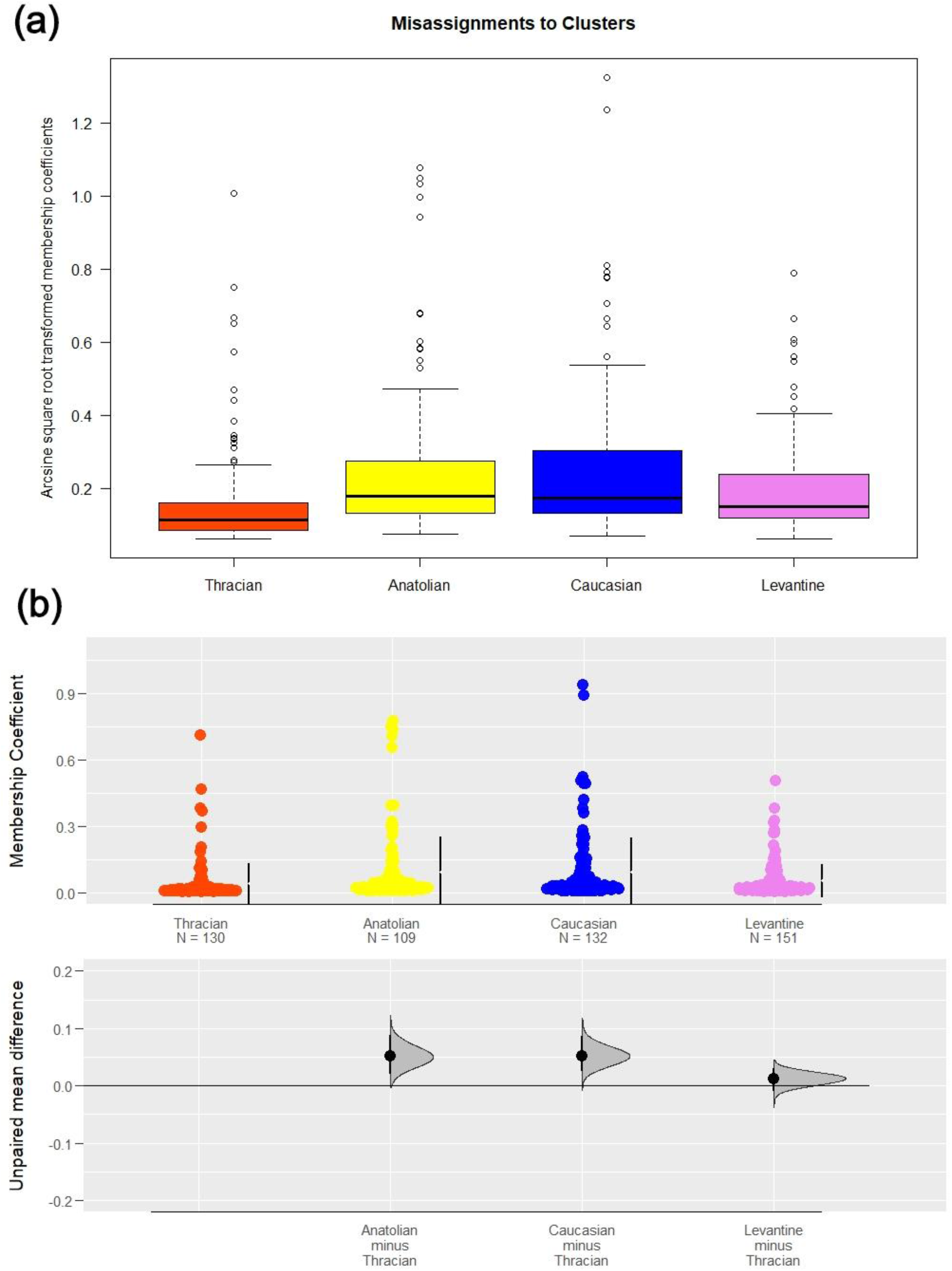
Comparison of misassignment proportions between the major clusters (a) boxplot display of arcsine root square transformed membership coefficients (b) scatter plot with estimations of mean differences based on raw individual membership coefficients (orange: Thracian, yellow: Anatolian, blue: Caucasian, violet: Levantine clusters).

A significant Kruskal-Wallis test (p < 0.001) and a *post hoc* Dunn’s test, accompanied by a significant ANOVA result (p <0.001) followed by a Tukey’s test, showed that misassignments to *A. m. caucasica* and *A. m. anatoliaca* clusters were significantly more frequent than the others (p <0.001 for both subspecies against C-lineage Thracian bees and p <0.05 against *syriaca* group). The effect sizes according to Cohen’s *d* varied from 0.34 to 0.54 with estimation plots verifying the precision of the difference observed (Fig. 6b). Despite observation of the highest values in *A. m. caucasica* misassignments, the results were not significant between *A. m. caucasica* and *A. m. anatoliaca* clusters.

We checked if those differences result from many individuals with high admixture levels but such data only constituted the 7.5% of all the observations. This is with a threshold level of 0.5 for transformed values which corresponds to a second hybrid with a 25% contribution of non-native origin. So, we concluded that rather the main effect is due to consistent mid to low subspecies’ contributions to other populations.

We also investigated if these small drifts in admixture proportions were more prominent in some localities and if populations differed in the subspecies they are receiving gene flow. This led us to comprehend the extent, magnitude and direction of the patterns of gene flow among the subspecies with a particular sensitivity to the populations. The results are summarized in Fig 7. We applied Dunn’s test for each pairwise comparison between the populations. Significance of results will be mentioned but details of the test results can be found in the supplementary file.

**Figure 7.**
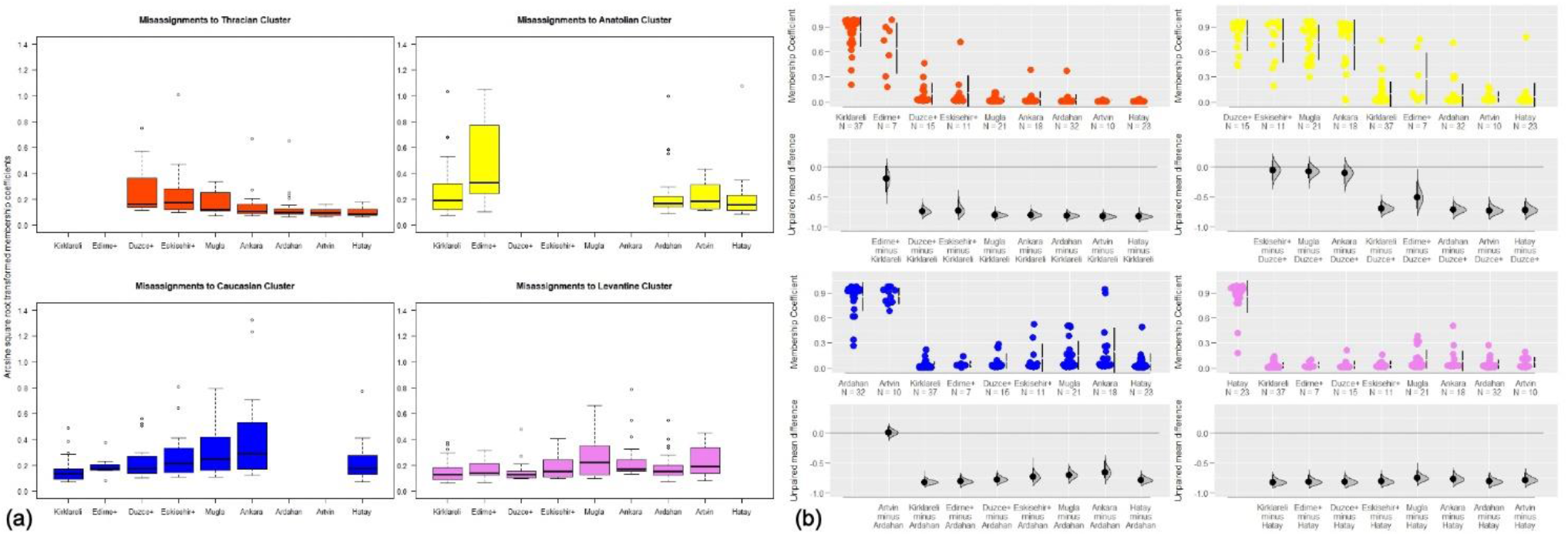
Patterns of gene flow between populations (a) boxplot display of arcsine root square transformed membership coefficients (b) scatter plot with estimations of mean differences based on raw individual membership coefficients (orange: Thracian, yellow: Anatolian, blue: Caucasian, violet: Levantine clusters).

Contributions from the Thracian cluster seem to be high in Düzce+ and Eskişehir+ which reside in the southeast of Marmara Sea across the Bosphorus and also there is some non-significant surplus in Muğla province in the Aegean coast. This is consistent with the Structure result for K = 2. While Thracian populations of Kırklareli and Edirne+ receive most gene flow from the Anatolian cluster these are not significantly different than Anatolian contributions to other regions which points to a balanced, uniform contribution of this subspecies to each group. Although non-significant, Edirne+ receives much gene flow from that cluster. Caucasian cluster on the other hand contributes most to Ankara, Muğla, Eskişehir+ and Düzce+ populations. Only significant differences are observed between Ankara and Muğla receiver populations and Kırklareli in terms of admixture with Caucasian populations. The same populations also significantly differed from Kırklareli in their admixture levels with the Levantine cluster. In both cases Kırklareli population in Thrace had lower gene flow. This is interesting since Muğla in the southwest and Caucasus region at the northeast lie at the different extremes of the country.

## Discussion

FST values obtained were highly significant but they were lower than what Bodur et al. (2007) estimated -a total F_ST_ of 0.077 together with higher values for pairwise comparisons among populations- by samples collected ten years before ours. This may indicate a recent increased gene flow and can be an alarm signal for a trend. Constant monitoring studies are needed in the future to see if it is a persistent trend really. The high degree of structuring in stationary colonies according to FST results was lost in migratory ones, meaning they are less differentiated from each other due to high degree of gene flow.

Phylogenetic tree clearly showed that Thracian samples were completely distinct from others pointing to an early division of populations and limited gene flow. This supports the hypothesis for a Carniolan (C-lineage) descent of Thracian bees in Turkey. Directly including samples from the major C-lineage subspecies would confirm the subspecies of these bees highly differentiated from Anatolian samples. In our initial observations of another research, Thracian samples grouped with C-lineage European breeds rather than the samples throughout Anatolia (Kükrer, unpublished data). This is a challenging point to Ruttner’s claim (Ruttner 1988) that Thracian bees belong to *anatoliaca* subspecies and needs further investigation.

West Anatolian, Levant and Caucasian populations did also form separate clusters in the tree. PCA results confirmed those 4 different clusters inferred from tree topology. Bitlis+ samples resided with Central and West Anatolian populations in both phylogenetic tree and PCA results (supplementary file) but it should be kept in mind that all samples from that locality belonged to migratory colonies so resampling with inclusion of stationary colonies from East Anatolia would be beneficial to understand the real phylogenetic relations.

The two most possible K values in structure analysis for the whole sample and the stationary colonies were K = 2 and K = 4, both results supporting the hypotheses of populations belonging to 2 separate lineages (C and O) and 4 distinct subspecies (a Carniolan ecotype in Thrace, *A. m. caucasica* in Artvin and Ardahan, *A. m. syriaca* in Hatay and *A. m. anatoliaca* widely distributed covering the rest of the country). In contrast to the expectations of migratory beekeepers of making use of native stocks, results involving migratory beekeepers’ samples lacked any population structuring in the cluster analysis further clarifying the highly hybridized status of migratory apiaries.

Stationary apiaries, as expected, yielded highly structured groups where all the subspecies could be detected. When K was 2, the structure analysis of two distinct clusters showed that there was a transition zone between Thracian and Anatolian samples around Marmara Sea and Aegean. This may be a hybrid zone between the C and O lineages like the ones identified before between M and C lineages in Alps and Apennine Peninsula and between A and M lineages at the Iberian Peninsula and Mediterranean islands (De la Rua et al. 2009). An analysis of ecological niches under species distribution models suggest an intersection of habitat suitability of both subspecies within the aforementioned geographic area (Kükrer, unpublished data).

When K was considered as 4, all four subspecies were easily differentiated from each other, in accordance with the expectances. The significance of two distinct clusters (K = 2) was higher than four (K = 4) which means that the differences between the populations belonging to C (in Thrace) and O (in Anatolia) lineages are more clear-cut than differences between the populations of four different subspecies.

*A. m. anatoliaca* samples fell in the middle of the other subspecies in ordinations, being similar to all other populations according to FST values despite being a distinct cluster in structure analysis which may point to a significant historical contribution to *A. m. anatoliaca* populations from the neighboring regions. Another explanation can be that *anatoliaca* subspecies’ putative basal position for O-lineage honey bees places it as a center of genetic diversity. With *anatoliaca* bees exhibiting a distinct identity, the situation was quite different than what was observed in all-migratory Bitlis+ samples where a mixture of different clusters surpassed instead of a separate identity.

A better understanding in terms of phylogenetic relationships between the populations in Turkey can be developed if populations neighboring Anatolia and Thrace in Balkans, Iran, Caucasus and the Middle East are also sampled. This can be a direction for future research, for shedding light on the complicated taxonomic status within and between the C and O lineages and for drawing edges and transition zones of the subspecies present across the whole region.

Results from different analyses conducted here confirmed the presence of clusters but also, they all together pointed to the status of migratory colonies: they might be acting as a hybrid zone mobile in space and time, being at one region in spring and at others in summer and fall, becoming vectors of otherwise local gene combinations. Statistical results concerning a comparison between migratory and stationary colonies confirmed the significant gene flow towards the migrants from local bees.

A significant gene flow towards local bees was also observed by a comparison between isolated regions and those are not. This result, derived from directly contrasting two settings in an experimental framework, is pointing to the vitality of establishing areas away and free from migratory beekeeping for preservation of honey bee genetic diversity in conclusion with other studies on conservation practices (Pinto et al. 2015; Oleksa et al. 2015).

One interesting point in the results was that the trend of the stationary colonies in Ankara. They had a significantly lower probability of being assigned to their own clusters than the migratory colonies of their province. This may be related with the regions migratory beekeepers of Ankara prefer to visit during their migratory cycle or due to the insistent preference of using native queen bees by migratory beekeepers. The low assignment degree of stationary colonies in Ankara may also be related with Kazan apiary of TKV (Development Foundation of Turkey) placed there where hundreds of colonies of Caucasian bees are raised and sold around for more than 30 years. The same practice is also carried out by many queen bee breeders in Kazan region. Gene flow through these apiaries and queen bees distributed locally by trade may contribute quietly to such an admixture observed in stationary colonies in Ankara. The high misassignment probability of colonies in Ankara to the Caucasian cluster also revealed such a process as probable.

It’s hard to directly quantify the effect of queen and colony trade but unique features of Anatolia and Thrace by availability of a number of naturally occurring subspecies renders possible the understanding of their relative roles. Honeybees from stationary colonies were assigned more often to their native clusters but they were also assigned to other clusters with lower probabilities. Samples in the whole range of the study misassigned to Caucasian cluster more often than they were misassigned to others.

This is most probably due to wide distribution of Caucasian queen bees by trade. Migratory beekeeping is not practiced in Ardahan and Artvin where highly commercial Caucasian bees are native. Hence no bees go in or leave out the region as migratory colonies. So, the observed introgression of Caucasian alleles to the stationary colonies elsewhere whose beekeepers let them change their queens on their own rather than purchasing queens of different origins, could mainly be attributed to frequent queen bee and colony replacements in neighboring apiaries within those regions.

It is shown here that practicing of honey bee replacements increase the level of admixture within the gene pool. As previously discussed, a very high level of Caucasian introgression was observed in Ankara. *A. m. anatoliaca* alleles also showed high introgression especially in Edirne+ of Thrace region but also at average levels in other regions. These high levels may be related to this subspecies’ geographical proximity to other populations which might have led to historical and recent gene exchange. By another explanation it can be related to the widespread practicing of migratory beekeeping by Western and Central Anatolian beekeepers throughout Turkey, rather than queen or colony replacements since there are very few commercial queen breeders within the distribution range of *A. m. anatoliaca*.

Results of the various statistical tests carried out and analysis applied in this study clearly showed that the genetic structure of honeybee populations in Turkey were highly conserved and still maintained. But it doesn’t mean that the structure and diversity observed is secure. Rather it should be considered under threat since the anthropogenic factors leading to gene flows are still underway and keep admixing the populations.

A quiet interesting point was that, the preservation of population structure was achieved despite a very high number of colonies moved from one location to the other by migratory beekeeping practice and despite unregulated and frequent queen and colony sales. Future research may also need to focus on how this biodiversity and its structuring were preserved and its relation to natural selection. Further hypothesis can be formulated to distinguish the relative effects of natural selection and gene flow, the former could be so significant that it could potentially counterbalance the latter.

Genetic variation eventually leading to local adaptations with such significant outweighing effect can be considered as a valuable resource for honey bee populations in the global context at this time of unusual bee losses as well as global climate change. So, a better understanding of both present adaptation to local climate and geographic conditions as well as adaptive capacity to future changes would better be developed for the sake of the bees and their beneficiaries. A fair amount of effort should be invested on more studies focusing on candidate functional variants at the genome level that play role in due process in different parts of the world. Novel and innovative ways of coping with environmental and climatic stressors developed by honey bee populations or exploration of interesting patterns of convergent evolution are waiting ahead to be yet discovered.

Importance of establishing isolated regions was highlighted with genetic data. The results of the statistical tests showed a significant difference between the conservation of identity in and out of isolated regions with isolated regions staying purer in terms of subspecies composition. Such regions were proven to be effective in conservation of unique diversity present within.

In the light of this study we propose a renewed effort to address the need for massive establishing of such regions for conserving locally adapted native bees throughout the whole natural distribution of the species. This especially holds for underrepresented regions in terms of local diversity hotspots. A gap analysis aiming for complementarity in the planning of systematic conservation efforts are urgently needed globally.

In such isolated regions, naturally, migratory beekeeping as well as replacement of queen bees with non-native ones must be strictly prohibited and checked by relevant molecular monitoring techniques. However, these isolated regions should also be wide enough involving additional buffer zones where further restrictions on migratory beekeeping and bee trade are applied for efficient isolation and for fulfilling sufficient effective population sizes.

Thanks to increasing awareness in the last decade within the industry, now there are at least 11 isolated regions in service or being established in Turkey through significant efforts of scientists and their collaboration with Turkish Beekeepers Association. There is an ever-growing need for establishing closer links with decision makers and stakeholders and necessity of investing more effort in communicating results of scientific studies in order to make the most out of them.

Queen bee trade is not currently subjected to any restrictions or regulations in Turkey and there are still very few pioneering measures within the natural distribution range, obviously not enough to guarantee the realized preservation in the next decades. Such measures should be applied from a conservation perspective to avoid extinction of native races, ecotypes and diversity present in these populations. Genetic similarity of donor and recipient populations should be considered while determining migration routes for migratory beekeepers and determining permissions for bee sales.

Central and western Anatolian populations suffer from significant gene flow from Caucasian populations as demonstrated by our results. Muğla and Ankara especially showed alarming levels of significant gene flow from other subspecies. This is not unexpected since the former receives millions of migratory colonies during the honeydew season.

Despite its wide range of distribution spanning Anatolia from one side to other, special consideration should be taken for preserving *A. m. anatoliaca* subspecies. The large and heterogenous native range of this subspecies permitted the evolution of numerous ecotypes including those in coastal, inner step or rainy forest ecosystems with noteworthy adaptations linked to their local environments.

The case with Hatay’s *syriaca* populations too, can get worse and worse since the migratory beekeeping practice is heavily carried out in the region and queen bee replacement with non-native races was frequent throughout the last decade. This is mainly due to aggressive behavior, high swarming tendency and an infame for low levels of honey piling but the subspecies is also shown to exhibit some natural forms of varroa resistance (Kence et al. 2013). In the future this may end up in *A. m. syriaca* colonies getting limited to a few localities and apiaries since the range of the subspecies in Turkey is very narrow. A long-term conservation program considering improvement of traits that result in beekeepers staying away from that subspecies should be actualized immediately in this region too.

Thracian populations show a significant differentiation from the rest of the bees in Anatolia but the subspecies which they belong to is not characterized on a strong basis yet and this unique population is not registered officially like the case with *A. m. syriaca* of Hatay. Only subspecies officially recognized in Turkey is *A. m. caucasica* so identification and registration procedures for the others should be put into practice as soon as possible.

An improvement based on molecular genetic techniques can be applied to the ongoing conservation programs for the *A. m. caucasica* subspecies. It is interesting to note that we even detected hybrid individuals within the range of largest, oldest and heavily invested conservation area. This proposal for application of molecular monitoring techniques holds for other subspecies too.

Recently a registration procedure for Muğla bees as an Aegean coastal ecotype of *anatoliaca* subspecies with specific adaptations to resource phenology in the form of availability of honeydew obtained from scale insect *M. hellenica* of Turkish red pine *P. brutia* is under process. During the conservation and breeding efforts, an adequate level of use of molecular markers was achieved (Kükrer, unpublished data). More attention should be paid to genetically characterize *A. m. meda* subspecies that was out of the reach of this study and which can be threatened by anthropogenic factors listed and studied here.

Rather than queen bee replacement, it should be encouraged to use native bees improved for desired characters which are also locally adapted by definition. Such improved breeds would be used locally and not distributed in a country-wide manner so that local adaptations would still be preserved while bees are selected for resistance to pests and pathogens, hygienic behavior, reduced aggressiveness, reduced tendency for swarming, higher winter survival, higher productivity or for increased pollination. For obtaining better results in that, research concerning the smoothing, development and extension of breeding locally adapted native bees and artificial insemination techniques should be given higher priority and be adopted globally throughout the natural distribution range of local subspecies.

Our overall results answer arguments about the present situation of honey bee subspecies in Turkey but they also bear a wider interest to the community since they constitute an important pioneer attempt to quantify the effects of human impact. Our measurable and justified scientific findings on migratory beekeeping, queen and colony trade as well as conservation implications will hopefully be of some use for the decision makers and other stakeholders.

## Supporting information

R codes

## Author Contributions

All authors conceived and planned the experiments and contributed to the field work. Mert Kükrer carried out the experiments and statistical analyses. All authors contributed to the interpretation of the results. Mert Kükrer took the lead in writing the manuscript. All authors provided critical feedback and helped shape the research, analysis and manuscript. Aykut Kence was in charge of overall direction and planning.

## Acknowledgments

We dedicate this paper to the memory of late Professor Aykut Kence who spent His last 20 years studying honey bee diversity in Turkey trying strongly to draw attention of researchers, decision makers and beekeepers to the conservation of locally adapted native bees as a precious legacy of our kind. He will be remembered also for defending theory of evolution to be taught in science curriculum and for training many valuable young evolutionary biologists under very harsh conditions. He passed away on Feb. 1. 2014 after the completion of the study and at the very beginning of manuscript preparation.

We’d like to thank numerous beekeepers who provided samples to the study and The Turkish Beekeepers Association. Special thanks to Dr. Devrim Oskay for his contribution in the form of consumables. We’d like to thank Mustafa Nail Cırık, Mehmet Ali Döke, Okan Can Arslan, Cansu Özge Tozkar, Mehmet Kayım and Eda Gazel Karakaş for the field work and Esin Öztürk and Ezgi Ersin as well as Batuhan Çağrı Yapan, Ayshin Ghalici, Gizem Kars and Batuhan Elçin for contributing lab sessions. Special thanks to Çiğdem Akın Pekşen and Cemal Can Bilgin for their advice on the early versions of the manuscript.

This manuscript has been released as a pre-print at https://www.biorxiv.org/content/10.1101/154195v1, (Kükrer et al. 2017) and includes elements from the corresponding authors thesis (Kükrer 2013).

## Supplementary material

R codes and Supplementary File.

