## Supplementary File for "Honey Bee Diversity is Swayed by Migratory Beekeeping and Trade Despite Conservation Practices: Genetic Evidences for the Impact of Anthropogenic Factors on Population Structure"

### *Supplementary Material*

#### 1 Supplementary Tables

Supplementary Table 1. Microsatellite loci and multiplex primer groups.

| | Locus | F/R | Sequence | Length | Label |
|---|---|---|---|---|---|
| | Ap218 | F | AGGGATGGAATTCTTCGATT | 20 | 6-FAM |
| | | R | TTGTCACAATTCCGCTTGA | 19 | |
| | A113 | F | CTCGAATCGTGGCGTCC | 17 | 6-FAM |
| | | R | CCTGTATTTTGCAACCTCGC | 20 | |
| | A(B)024 | F | CACAAGTTCCAACAATGC | 18 | VIC |
| | | R | CACATTGAGGATGAGCG | 17 | |
| GR 1 | Ap249 | F | CGCGCGACGACGAAATGT | 18 | VIC |
| | | R | CAGTCCTTTGATTCGCGCTACC | 22 | |
| | A088 | F | CGAATTAACCGATTTGTCG | 19 | NED |
| | | R | GATCGCAATTATTGAAGGAG | 20 | |
| | AP001 | F | ACACGCGAACAATACAACA | 19 | NED |
| | | R | ACTAATCGGCACGATGAAG | 19 | |
| | Ap043 | F | GGCGTGCACAGCTTATTCC | 19 | PET |
| | | R | CGAAGGTGGTTTCAGGCC | 18 | |
| | A079 | F | CGAAGGTTGCGGAGTCCTC | 19 | 6-FAM |
| | | R | GTCGTCGGACCGATGCG | 17 | |

| | | | | | |
|---|---|---|---|---|---|
| | Ac306 | F | GAATATGCCGCTGCCACC | 18 | 6-FAM |
| | | R | TTTCGTTGCATCCGAGCG | 18 | |
| | Ap226 | F | AACGGTGTTCGCGAAACG | 18 | 6-FAM |
| | | R | AGCCAACTCGTGCGGTCA | 18 | |
| **GR 2** | A007 | F | CCCTTCCTCTTTCATCTTCC | 20 | VIC |
| | | R | GTTAGTGCCCTCCTCTTGC | 19 | |
| | HB-C16-01 | F | AAAATGCGATTCTAATCTGG | 20 | VIC |
| | | R | TTGCCTAAAATGCTTGCTAT | 20 | |
| | Ap068 | F | TGTCTGCCCTCCTCTCTGTT | 20 | NED |
| | | R | CACATCGAGCGAGAAGGC | 18 | |
| | A014 | F | GTGTCGCAATCGACGTAACC | 20 | NED |
| | | R | GTCGATTACCGATCGTGACG | 20 | |
| | Ap223 | F | TCGTACAACGTCGCGCAA | 18 | PET |
| | | R | GCCGCTCGCCTGTATCTG | 18 | |

Supplementary Table 1. Cont. Microsatellite loci and multiplex primer groups.

| | Locus | F/R | Sequence | Length | Label |
|---|---|---|---|---|---|
| | AP019 | F | CTCGTTTCTTCCATTGCG | 18 | 6-FAM |
| | | R | CGGTACGCGGTAGAAAGA | 18 | |
| | A(B)124 | F | GCAACAGGTCGGGTTAGAG | 19 | 6-FAM |
| | | R | CAGGATAGGGTAGGTAAGCAG | 21 | |
| | A043 | F | CACCGAAACAAGATGCAAG | 19 | VIC |
| | | R | CCGCTCATTAAGATATCCG | 19 | |
| **GR 3** | A076 | F | GCCAATACTCTCGAACAATG | 20 | VIC |
| | | R | GTCCAATTCACATGTCGACATC | 22 | |
| | Ap273 | F | GATCTTGTGTTAAACAGCCG | 20 | NED |
| | | R | GATCTCTGGCAGACGAAGAG | 20 | |
| | Ap289 | F | AGCTAGGTCTTTCTAAGAGTGTTG | 24 | NED |
| | | R | TTCGACCGCAATAACATTC | 19 | |
| | HB-C16-05 | F | ATTTTATGCGCGTTTCGTA | 19 | PET |
| | | R | CATGGCTCCTCCATTAAATC | 20 | |
| | A028 | F | GAAGAGCGTTGGTTGCAGG | 19 | PET |
| | | R | GCCGTTCATGGTTACCACG | 19 | |
| | Ap049 | F | CCAATAGCGGCGAGTGTG | 18 | 6-FAM |

| | | | | | |
|---|---|---|---|---|---|
| | | R | GGGCTTCGTACGTCCACC | 18 | |
| | Ap238 | F | GTCTCGTGCGTGCGAATG | 18 | 6-FAM |
| | | R | TTCATCATGTTCTCAAATTTCTTTGT | 26 | |
| | AC006 | F | GATCGTGGAAACCGCGAC | 18 | VIC |
| | | R | CACGGCCTCGTAACGGTC | 18 | |
| **GR 4** | Ap243 | F | AATGTCCGCGAGCATCTG | 18 | VIC |
| | | R | TGTTTACGAGAATTCGACGGG | 21 | |
| | Ap288 | F | GTTAGTTCGTCGTCGACCG | 19 | NED |
| | | R | TCTTAGCTTTATAACGAGCACG | 22 | |
| | HB-C16-02 | F | TAGTATCGTGCTGTTCATCG | 20 | NED |
| | | R | ACATACATCTCTTGGCGAGT | 20 | |
| | A107 | F | CCGTGGGAGGTTTATTGTCG | 20 | PET |
| | | R | CCTTCGTAACGGATGACACC | 20 | |

Supplementary Table 2. PCR conditions.

| | STEP | TIME | TEMPERATURE |
|---|---|---|---|
| Activation | | 5 minutes | 95 ºC |
| 30 cycles | Denaturation | 30 seconds | 95 ºC |
| | Annealing | 150 seconds | 57 ºC |
| | Extension | 30 seconds | 72 ºC |
| Final extension | | 30 minutes | 60 ºC |

Supplementary Table 3. Null allele frequencies.

| LOCUS | POPULATION | FREQUENCY |
|---|---|---|
| A(B)024 | Artvin | 0.20410 |
| Ap273 | Eskişehir+ | 0.21365 |
| Ap289 | Ardahan | 0.36818 |
| Ap289 | Artvin | 0.40463 |

Supplementary Table 4. Private alleles for populations.

| POPULATION | LOCUS | ALLELE | FREQUENCY |
|---|---|---|---|
| Kırklareli | AB124 | 213 | 0,053 |
| Edirne+ | A007 | 124 | 0,063 |
| Edirne+ | A007 | 149 | 0,063 |
| Edirne+ | AB124 | 244 | 0,063 |
| Edirne+ | AP001 | 252 | 0,063 |
| Edirne+ | AP223 | 182 | 0,125 |
| Düzce+ | AP238 | 262 | 0,067 |
| Muğla | AP289 | 200 | 0,056 |
| Artvin | A079 | 127 | 0,050 |
| Artvin | A107 | 185 | 0,050 |
| Artvin | A113 | 242 | 0,050 |
| Artvin | AP249 | 216 | 0,050 |
| Hatay | AP243 | 268 | 0,056 |

Supplementary Table 5. Loci based allelic diversity.

| LOCI | # of ALLELES | # of EFFECT. ALLELES | # of PRIVATE ALLELES | ALLELIC RICHNESS | INFORM. INDEX | HOMOZY. OBS. |
|---|---|---|---|---|---|---|
| A007 | 46 | 19,0 | 11 | 11,6 | 3,3 | 0,05 |
| A014 | 9 | 2,2 | 3 | 3,2 | 1,0 | 0,45 |
| A028 | 8 | 1,3 | 4 | 2,3 | 0,5 | 0,75 |
| A043 | 13 | 2,0 | 5 | 3,8 | 1,1 | 0,50 |
| A079 | 12 | 3,9 | 4 | 4,9 | 1,6 | 0,26 |
| A088 | 13 | 2,8 | 4 | 3,9 | 1,3 | 0,36 |
| A107 | 25 | 13,6 | 2 | 10,5 | 2,9 | **0,07** |
| A113 | 23 | 9,3 | 7 | 8,3 | 2,4 | 0,11 |
| A(B)024 | 8 | 2,5 | 2 | 3,5 | 1,1 | 0,39 |
| A(B)124 | 16 | 4,8 | 4 | 6,4 | 2,0 | 0,21 |
| AC006 | 8 | 1,2 | 2 | 2,2 | 0,5 | 0,80 |
| AC306 | 11 | 3,5 | 3 | 4,3 | 1,4 | 0,29 |
| AP001 | 33 | 4,7 | 7 | 7,1 | 2,2 | 0,21 |
| AP019 | 8 | 1,6 | 2 | 2,9 | 0,8 | 0,64 |
| AP043 | 33 | 7,6 | 6 | 8,5 | 2,5 | 0,13 |
| AP049 | 13 | 1,9 | 4 | 3,9 | 1,1 | 0,52 |
| AP068 | 9 | 2,8 | 2 | 4,5 | 1,4 | 0,35 |
| AP218 | 6 | 1,1 | 3 | 1,8 | 0,3 | 0,87 |
| AP223 | 6 | 3,1 | 1 | 3,9 | 1,3 | 0,33 |
| AP226 | 7 | 1,3 | 3 | 2,4 | 0,5 | 0,79 |
| AP238 | 6 | 1,8 | 3 | 2,3 | 0,7 | 0,57 |
| AP243 | 11 | 1,2 | 6 | 2,4 | 0,5 | 0,81 |
| AP249 | 11 | 3,6 | 1 | 5,2 | 1,6 | 0,28 |
| AP273 | 4 | 1,8 | 1 | 2,1 | 0,7 | 0,56 |
| AP288 | 7 | 1,7 | 2 | 3,2 | 0,8 | 0,59 |
| AP289 | 40 | 10,7 | 5 | 9,9 | 2,9 | 0,09 |
| HB-C16-01 | 40 | 16,8 | 2 | 11,5 | 3,2 | 0,06 |
| HB-C16-02 | 35 | 3,7 | 10 | 7,2 | 2,2 | 0,27 |
| HB-C16-05 | 5 | 2,8 | 1 | 3,1 | 1,1 | 0,36 |
| MEAN | 16,1 | 4,6 | 3,8 | 5,1 | 1,5 | 0,40 |

Supplementary Table 6.. Abundances of alleles per population.

| | # of ALLELES | # of FREQ. ALLELES | # of EFFECT. ALLELES | # of PRIVATE ALLELES | # of LOCAL COMMON ALLELES |
|---|---|---|---|---|---|
| **KIRKLARELİ** | 255 | 115 | 113 | 19 | 31 |
| **EDİRNE+** | 135 | 135 | 96 | 5 | 16 |
| **DÜZCE+** | 163 | 103 | 92 | 6 | 17 |
| **ESKİŞEHİR+** | 140 | 89 | 85 | 5 | 10 |
| **ANKARA** | 236 | 102 | 117 | 11 | 24 |
| **MUĞLA** | 245 | 102 | 123 | 14 | 27 |
| **ARDAHAN** | 209 | 89 | 101 | 13 | 19 |
| **ARTVİN** | 138 | 138 | 87 | 4 | 15 |
| **BİTLİS+** | 270 | 103 | 126 | 7 | 35 |
| **HATAY** | 287 | 101 | 130 | 26 | 32 |
| **MEAN** | 207,8 | 107,7 | 106,9 | 11,0 | 22,6 |

Supplementary Table 7. Allelic diversities per population.

| | INFO. INDEX | GENE DIVERSITY | ALLELIC RICHNESS |
|---|---|---|---|
| **KIRKLARELİ** | 1,4 | 0,62 | 4,9 |
| **EDİRNE+** | 1,2 | 0,62 | 4,7 |
| **DÜZCE+** | 1,2 | 0,58 | 4,4 |
| **ESKİŞEHİR+** | 1,1 | 0,55 | 4,3 |
| **MUĞLA** | 1,3 | 0,55 | 4,6 |
| **ANKARA** | 1,3 | 0,58 | 4,8 |
| **ARDAHAN** | 1,1 | 0,50 | 4,2 |
| **ARTVİN** | 1,0 | 0,51 | 4,3 |
| **BİTLİS+** | 1,4 | 0,59 | 4,9 |
| **HATAY** | 1,4 | 0,60 | 5,1 |
| **MEAN** | 1,2 | 0,57 | 4,6 |

Supplementary Table 8. Observed and expected heterozygosities for each loci.

| Locus | Obs. Het. | Exp.Het. |
|---|---|---|
| A007 | 0,90 | 0,95 |
| A014 | 0,46 | 0,55 |
| A028 | 0,24 | 0,25 |
| A043 | 0,39 | 0,50 |
| A079 | **0,69** | 0,74 |
| A088 | 0,58 | 0,64 |
| A107 | **0,92** | 0,93 |
| A113 | 0,82 | 0,89 |
| A(B)024 | 0,56 | 0,61 |
| A(B)124 | **0,73** | 0,79 |
| AC006 | **0,15** | 0,20 |
| AC306 | **0,64** | 0,71 |
| AP001 | 0,68 | 0,79 |
| AP019 | 0,31 | 0,36 |
| AP043 | 0,79 | 0,87 |
| AP049 | 0,43 | 0,48 |
| AP068 | 0,70 | 0,65 |
| AP218 | **0,06** | 0,13 |
| AP223 | 0,64 | 0,68 |
| AP226 | **0,17** | 0,21 |
| AP238 | 0,43 | 0,43 |
| AP243 | **0,16** | 0,19 |
| AP249 | 0,70 | 0,72 |
| AP273 | 0,38 | 0,44 |
| AP288 | 0,36 | 0,41 |
| AP289 | **0,66** | 0,91 |
| HB-C16-01 | 0,88 | 0,94 |
| HB-C16-02 | **0,59** | 0,73 |
| HB-C16-05 | 0,62 | 0,64 |

Supplementary Table 9. Loci in linkage disequilibrium.

| POPULATION | LOCI |
|---|---|
| KIRKLARELİ | 1 & 12 |
| KIRKLARELİ | 16 & 22 |
| BİTLİS+ | 27 & 29 |
| HATAY | 7 & 27 |
| HATAY | 15 & 20 |
| HATAY | 19 & 20 |

Supplementary Table 10. Estimated effective population sizes.

| POPULATION | EFFECTIVE SIZE |
|---|---|
| THRACE | 1860.3 |
| WEST ANATOLIA | 3500.0 |
| NORTH EAST | 776.6 |
| BİTLİS+ | 4655.9 |
| HATAY | 665.0 |

#### 2 Supplementary Figures

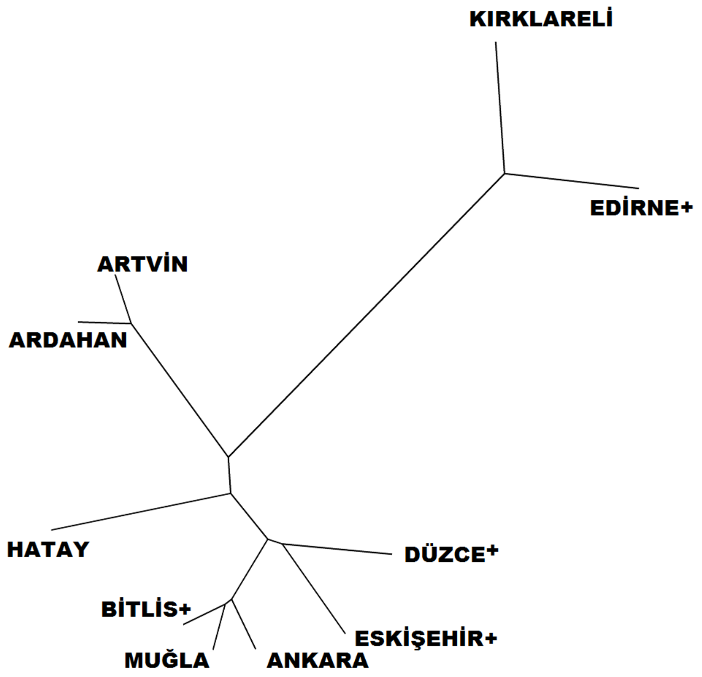


**Supplementary Figure 1.** UPGMA tree based on Nei's genetic distance (1978).

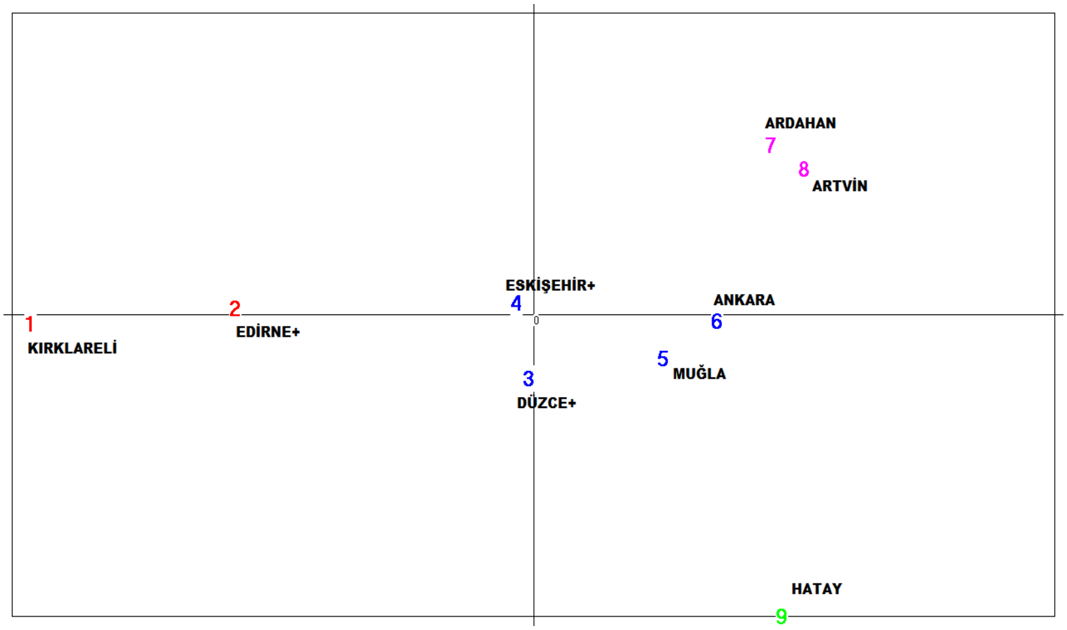


**Supplementary Figure 2.** PCA of stationary colonies. Axis 1: 43,19%. Axis 2: 21,87%.

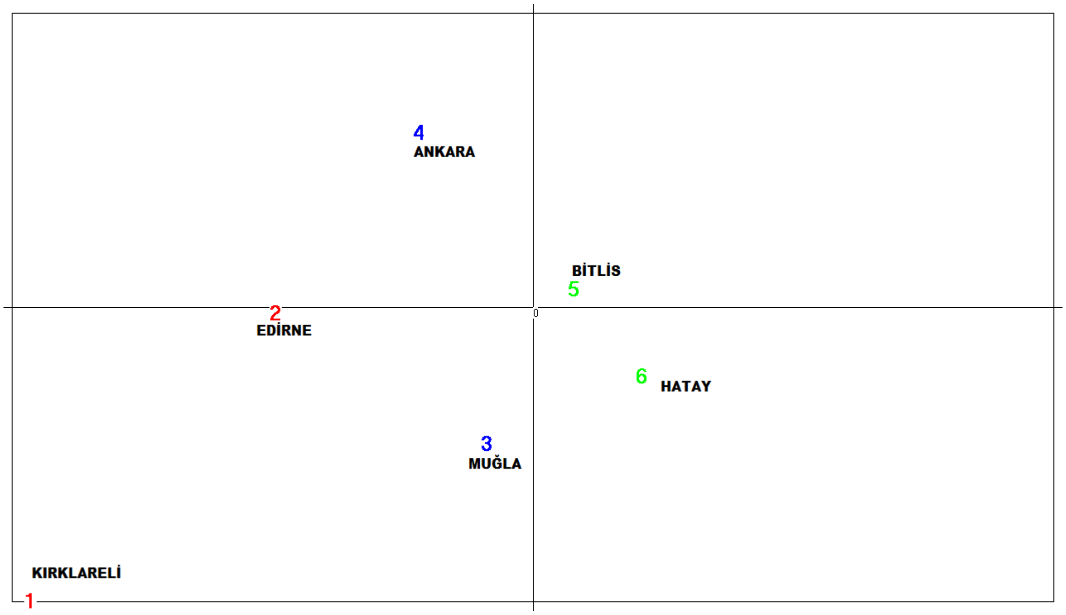


**Supplementary Figure 3.** PCA of migratory colonies. Axis 1: 28,21%. Axis 2: 20,98%.

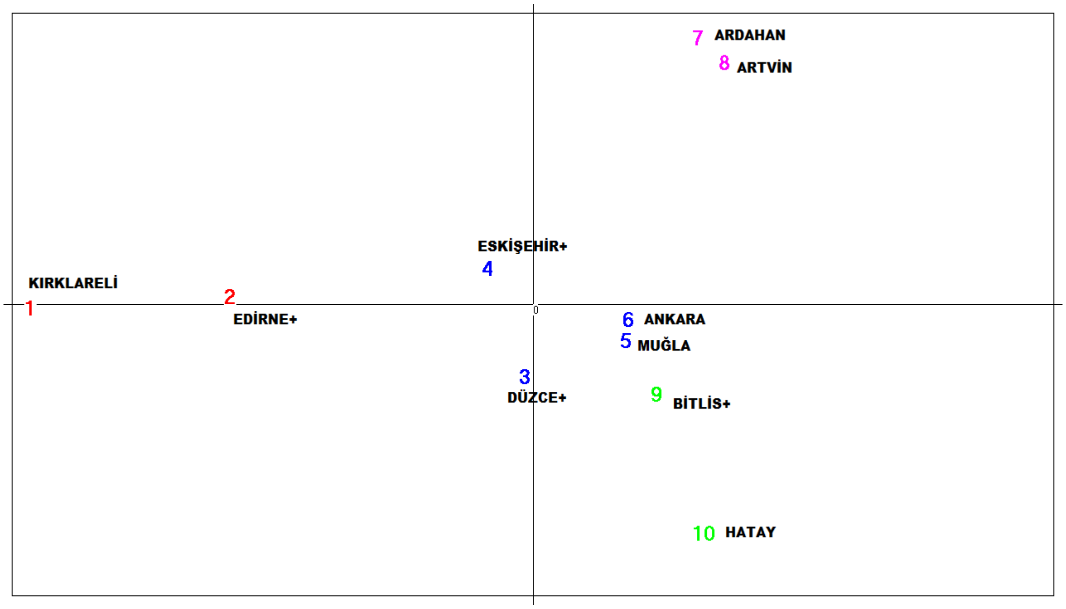


**Supplementary Figure 4.** PCA of overall data. Axis 1: 43,16%. Axis 2: 22,12%.

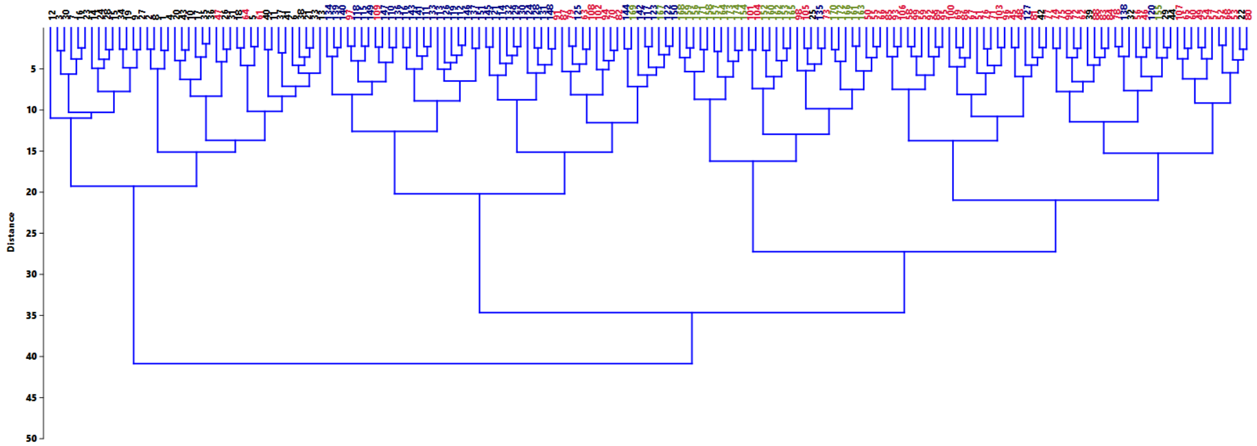

**Supplementary Figure 5.** Phylogenetic tree constructed using Ward's method and Euclidean distances (black: Thracian, blue: Caucasian, olive: Levantine, red: Anatolian clusters )
