## Supplementary material for "Honey Bee Diversity is Swayed by Migratory Beekeeping and Trade Despite Conservation Practices: Genetic Evidences for the Impact of Anthropogenic Factors on Population Structure": R codes

### Comparison

Mert Kukrer

27.04.2020

#### Table of Contents

##### Data

```
getwd()

## [1] "D:/Google Drive/UNIVERSITE/Akademik/Frontiers in ecology and evolutio
n/Latest version/Comparisons"

#install.packages(c("dunn.test","pwr", "effsize","dabestr"), type='binary', r
epos = "http://cran.rstudio.com/", dependencies = TRUE)

library(pwr)
library(effsize)
library(dunn.test)
library(dabestr)

## Loading required package: boot

## Loading required package: magrittr

source("D:/Google Drive/UNIVERSITE/Akademik/Frontiers in ecology and evolutio
n/Latest version/Comparisons/cbind.na.R")

memberships250 =read.csv2("D:/Google Drive/UNIVERSITE/Akademik/Frontiers in e
cology and evolution/Latest version/Comparisons/250_membership_coefficients.c
sv", header = TRUE, sep = ";", quote = "", dec = ",", fill = TRUE, comment.ch
ar = "")
head(memberships250)

##   SAMPLE_ID RANK POPULATION MIGRATORY THRACIAN LEVANTINE ANATOLIAN CAUCASI
AN
## 1      TK01    1 Kirklareli         0   0.8920    0.0450    0.0473    0.01
57
## 2      TK02    2 Kirklareli         0   0.9317    0.0137    0.0397    0.01
50
## 3      TK03    3 Kirklareli         0   0.9700    0.0090    0.0093    0.01
17
## 4      TK04    4 Kirklareli         0   0.8946    0.0280    0.0297    0.04
77
## 5      TK05    5 Kirklareli         0   0.8784    0.0210    0.0710    0.02
97
```

```
## 6      TK06    6 Kirklareli         0   0.8850    0.0247    0.0307    0.05
97
##   TransformedTHRACIAN TransformedLEVANTINE TransformedANATOLIAN
## 1            1.235940           0.21375613           0.21923770
## 2            1.306383           0.11731592           0.20059108
## 3            1.396713           0.09501121           0.09658661
## 4            1.240151           0.16812289           0.17320155
## 5            1.214600           0.14542582           0.26971656
## 6            1.224819           0.15781662           0.17612328
##   TransformedCAUCASIAN
## 1            0.1256298
## 2            0.1227828
## 3            0.1083786
## 4            0.2201780
## 5            0.1732016
## 6            0.2468347
```

```
memberships174 =read.csv2("D:/Google Drive/UNIVERSITE/Akademik/Frontiers in e
cology and evolution/Latest version/Comparisons/174_membership_coefficients.c
sv", header = TRUE, sep = ";", quote = "", dec = ",", fill = TRUE, comment.ch
ar = "")
head(memberships174)
```

```
##   SAMPLE_ID RANK POPULATION ANATOLIAN THRACIAN CAUCASIAN LEVANTINE
## 1      TK01    1 Kirklareli    0.0244   0.9415    0.0072    0.0269
## 2      TK02    2 Kirklareli    0.0292   0.9585    0.0057    0.0067
## 3      TK03    3 Kirklareli    0.0057   0.9835    0.0063    0.0045
## 4      TK04    4 Kirklareli    0.0284   0.9282    0.0292    0.0142
## 5      TK05    5 Kirklareli    0.0363   0.9404    0.0151    0.0082
## 6      TK06    6 Kirklareli    0.0243   0.9315    0.0267    0.0176
##   TransformedANATOLIAN TransformedTHRACIAN TransformedCAUCASIAN
## 1           0.15684730            1.326506           0.08495497
## 2           0.17172281            1.365645           0.07557025
## 3           0.07557025            1.441988           0.07945612
## 4           0.16933104            1.299526           0.17172281
## 5           0.19169752            1.324173           0.12319343
## 6           0.15652291            1.305987           0.16413736
##   TransformedLEVANTINE
## 1           0.16475656
## 2           0.08194521
## 3           0.06713245
## 4           0.11944759
## 5           0.09067807
## 6           0.13305726
```

#### Migratory beekeeping

```
Ankara_mig = subset(memberships250, memberships250$POPULATION=="Ankara")
Mugla_mig = subset(memberships250, memberships250$POPULATION=="Mugla")
Hatay_mig = subset(memberships250, memberships250$POPULATION=="Hatay")
Kirklareli_mig = subset(memberships250, memberships250$POPULATION=="Kirklarel
```

```
i")
Edirne_mig = subset(memberships250, memberships250$POPULATION=="Edirne+")
Duzce_mig = subset(memberships250, memberships250$POPULATION=="Duzce+")
Eskisehir_mig = subset(memberships250, memberships250$POPULATION=="Eskisehir+
")
Ardahan_mig = subset(memberships250, memberships250$POPULATION=="Ardahan")
Artvin_mig = subset(memberships250, memberships250$POPULATION=="Artvin")
Bitlis_mig = subset(memberships250, memberships250$POPULATION=="Bitlis+")

three_provinces_mig = cbind(
  rbind(
    Ankara_mig,
    Mugla_mig,
    Hatay_mig),
  TransformedCOEFFICIENT=c(
    Ankara_mig$TransformedANATOLIAN,
    Mugla_mig$TransformedANATOLIAN,
    Hatay_mig$TransformedLEVANTINE)
)

all_provinces_mig = cbind(
  rbind(
    Kirklareli_mig,
    Edirne_mig,
    Duzce_mig,
    Eskisehir_mig,
    Ankara_mig,
    Mugla_mig,
    Ardahan_mig,
    Artvin_mig,
    Bitlis_mig,
    Hatay_mig),
  TransformedCOEFFICIENT=c(
    Kirklareli_mig$TransformedTHRACIAN,
    Edirne_mig$TransformedTHRACIAN,
    Duzce_mig$TransformedANATOLIAN,
    Eskisehir_mig$TransformedANATOLIAN,
    Ankara_mig$TransformedANATOLIAN,
    Mugla_mig$TransformedANATOLIAN,
    Ardahan_mig$TransformedCAUCASIAN,
    Artvin_mig$TransformedCAUCASIAN,
    Bitlis_mig$TransformedLEVANTINE,
    Hatay_mig$TransformedLEVANTINE)
)

normalities_250 = apply(memberships250[5:12], 2, function(x) {shapiro.test(x)
$p.value})
p.adjust(normalities_250, method="BH")
```

```
##             THRACIAN             LEVANTINE             ANATOLIAN
##         6.204134e-22          8.026862e-20          4.700821e-15
##            CAUCASIAN   TransformedTHRACIAN TransformedLEVANTINE
##         3.016805e-18          7.246785e-20          7.819709e-17
## TransformedANATOLIAN TransformedCAUCASIAN
##         3.925518e-12          1.824682e-15
```

```
boxplot(
  Ankara_mig$TransformedANATOLIAN[Ankara_mig$MIGRATORY==0],
  Ankara_mig$TransformedANATOLIAN[Ankara_mig$MIGRATORY==1],
  Mugla_mig$TransformedANATOLIAN[Mugla_mig$MIGRATORY==0],
  Mugla_mig$TransformedANATOLIAN[Mugla_mig$MIGRATORY==1],
  Hatay_mig$TransformedLEVANTINE[Hatay_mig$MIGRATORY==0],
  Hatay_mig$TransformedLEVANTINE[Hatay_mig$MIGRATORY==1],
  three_provinces_mig$TransformedCOEFFICIENT[three_provinces_mig$MIGRATORY==0
],
  three_provinces_mig$TransformedCOEFFICIENT[three_provinces_mig$MIGRATORY==1
],
  all_provinces_mig$TransformedCOEFFICIENT[all_provinces_mig$MIGRATORY==0],
  all_provinces_mig$TransformedCOEFFICIENT[all_provinces_mig$MIGRATORY==1],
  col= rep(c("yellow", "yellow", "violet", "coral4", "firebrick"), each= 2),
  names= paste0(rep(c("Sta_", "Mig_"), times= 5), rep(c("Ankara", "Mugla", "H
atay", "Combine", "All"), each= 2)),
  las=2,
  xlab="",
  ylab="Arcsine square root transformed membership coefficients",
  pars=list(par(mar=c(8,5,4,2))),
  main="Stationary vs Migratory Colonies",
  at =c(1,2, 4,5, 7,8, 10,11, 13,14)
)

mean_mig = apply(cbind.na(
  Ankara_mig$TransformedANATOLIAN[Ankara_mig$MIGRATORY==0],
  Ankara_mig$TransformedANATOLIAN[Ankara_mig$MIGRATORY==1],
  Mugla_mig$TransformedANATOLIAN[Mugla_mig$MIGRATORY==0],
  Mugla_mig$TransformedANATOLIAN[Mugla_mig$MIGRATORY==1],
  Hatay_mig$TransformedLEVANTINE[Hatay_mig$MIGRATORY==0],
  Hatay_mig$TransformedLEVANTINE[Hatay_mig$MIGRATORY==1],
  three_provinces_mig$TransformedCOEFFICIENT[three_provinces_mig$MIGRATORY==0
],
  three_provinces_mig$TransformedCOEFFICIENT[three_provinces_mig$MIGRATORY==1
],
  all_provinces_mig$TransformedCOEFFICIENT[all_provinces_mig$MIGRATORY==0],
  all_provinces_mig$TransformedCOEFFICIENT[all_provinces_mig$MIGRATORY==1]),
  2, function(x) {mean(x, na.rm=TRUE)})
mean_mig
```

```
##  [1] 0.8237470 1.1147220 0.9256943 0.7033353 1.2027229 0.6633430 0.9988654
##  [8] 0.7893510 1.0590524 0.7158815
```

```r
effect_sizes_mig = c(
  cohen.d(TransformedANATOLIAN ~ MIGRATORY, data= Ankara_mig)$estimate,
  cohen.d(TransformedANATOLIAN ~ MIGRATORY, data= Mugla_mig)$estimate,
  cohen.d(TransformedLEVANTINE ~ MIGRATORY, data= Hatay_mig)$estimate,
  cohen.d(TransformedCOEFFICIENT ~ MIGRATORY, data= three_provinces_mig)$esti
mate,
  cohen.d(TransformedCOEFFICIENT ~ MIGRATORY, data= all_provinces_mig)$estima
te
)

## Warning in cohen.d.formula(TransformedANATOLIAN ~ MIGRATORY, data = Ankara
_mig):
## Cohercing rhs of formula to factor

## Warning in cohen.d.formula(TransformedANATOLIAN ~ MIGRATORY, data = Mugla_
mig):
## Cohercing rhs of formula to factor

## Warning in cohen.d.formula(TransformedLEVANTINE ~ MIGRATORY, data = Hatay_
mig):
## Cohercing rhs of formula to factor

## Warning in cohen.d.formula(TransformedCOEFFICIENT ~ MIGRATORY, data =
## three_provinces_mig): Cohercing rhs of formula to factor

## Warning in cohen.d.formula(TransformedCOEFFICIENT ~ MIGRATORY, data =
## all_provinces_mig): Cohercing rhs of formula to factor

effect_sizes_mig

## [1] -0.9987945  0.8859429  2.0124767  0.6661108  1.2218047

U_tests_mig = c(
  wilcox.test(TransformedANATOLIAN ~ MIGRATORY, data = Ankara_mig)$p.value,
  wilcox.test(TransformedANATOLIAN ~ MIGRATORY, data = Mugla_mig)$p.value,
  wilcox.test(TransformedLEVANTINE ~ MIGRATORY, data = Hatay_mig)$p.value,
  wilcox.test(TransformedCOEFFICIENT ~ MIGRATORY, data = three_provinces_mig)
$p.value,
  wilcox.test(TransformedCOEFFICIENT ~ MIGRATORY, data = all_provinces_mig)$p
.value
)
p.adjust(U_tests_mig, method="BH")

## [1] 1.317416e-02 1.472141e-02 1.718698e-04 4.750405e-03 1.280925e-12

F_tests_mig = c(
  var.test(TransformedANATOLIAN ~ MIGRATORY, data = Ankara_mig)$p.value,
  var.test(TransformedANATOLIAN ~ MIGRATORY, data = Mugla_mig)$p.value,
  var.test(TransformedLEVANTINE ~ MIGRATORY, data = Hatay_mig)$p.value,
  var.test(TransformedCOEFFICIENT ~ MIGRATORY, data = three_provinces_mig)$p.
value,
  var.test(TransformedCOEFFICIENT ~ MIGRATORY, data = all_provinces_mig)$p.va
```

```
lue
)
p.adjust(F_tests_mig, method="BH")
```

```
## [1] 0.25642361 0.54661092 0.01052260 0.29366739 0.01691253
```

```
t_tests_mig = c(
  t.test(TransformedANATOLIAN ~ MIGRATORY, data = Ankara_mig, var.equal = FAL
SE)$p.value,
  t.test(TransformedANATOLIAN ~ MIGRATORY, data = Mugla_mig, var.equal = FALS
E)$p.value,
  t.test(TransformedLEVANTINE ~ MIGRATORY, data = Hatay_mig, var.equal = FALS
E)$p.value,
  t.test(TransformedCOEFFICIENT ~ MIGRATORY, data = three_provinces_mig, var.
equal = FALSE)$p.value,
  t.test(TransformedCOEFFICIENT ~ MIGRATORY, data = all_provinces_mig, var.eq
ual = FALSE)$p.value
)
p.adjust(t_tests_mig, method="BH")
```

```
## [1] 1.054398e-02 1.634470e-02 5.627352e-04 5.172127e-03 4.000278e-12
```

```
pwr_t_tests_mig = c(
  pwr.t2n.test(
    n1= sum(Ankara_mig$MIGRATORY),
    n2= length(Ankara_mig$MIGRATORY) - sum(Ankara_mig$MIGRATORY),
    d= cohen.d(TransformedANATOLIAN ~ MIGRATORY, data= Ankara_mig)$estimate,
    sig.level= max(p.adjust(t_tests_mig, method="BH")))$power,
  pwr.t2n.test(
    n1= sum(Mugla_mig$MIGRATORY),
    n2= length(Mugla_mig$MIGRATORY) - sum(Mugla_mig$MIGRATORY),
    d= cohen.d(TransformedANATOLIAN ~ MIGRATORY, data= Mugla_mig)$estimate,
    sig.level= max(p.adjust(t_tests_mig, method="BH")))$power,
  pwr.t2n.test(
    n1= sum(Hatay_mig$MIGRATORY),
    n2= length(Hatay_mig$MIGRATORY) - sum(Hatay_mig$MIGRATORY),
    d= cohen.d(TransformedLEVANTINE ~ MIGRATORY, data= Hatay_mig)$estimate,
    sig.level= max(p.adjust(t_tests_mig, method="BH")))$power,
  pwr.t2n.test(
    n1= sum(three_provinces_mig$MIGRATORY),
    n2= length(three_provinces_mig$MIGRATORY) - sum(three_provinces_mig$MIGRA
TORY),
    d= cohen.d(TransformedCOEFFICIENT ~ MIGRATORY, data= three_provinces_mig)
$estimate,
    sig.level= max(p.adjust(t_tests_mig, method="BH")))$power,
  pwr.t2n.test(
    n1= sum(all_provinces_mig$MIGRATORY),
    n2= length(all_provinces_mig$MIGRATORY) - sum(all_provinces_mig$MIGRATORY
),
    d= cohen.d(TransformedCOEFFICIENT ~ MIGRATORY, data= all_provinces_mig)$e
stimate,
```

```
    sig.level= max(p.adjust(t_tests_mig, method="BH")))$power
)

## Warning in cohen.d.formula(TransformedANATOLIAN ~ MIGRATORY, data = Ankara
_mig):
## Cohercing rhs of formula to factor

## Warning in cohen.d.formula(TransformedANATOLIAN ~ MIGRATORY, data = Mugla_
mig):
## Cohercing rhs of formula to factor

## Warning in cohen.d.formula(TransformedLEVANTINE ~ MIGRATORY, data = Hatay_
mig):
## Cohercing rhs of formula to factor

## Warning in cohen.d.formula(TransformedCOEFFICIENT ~ MIGRATORY, data =
## three_provinces_mig): Cohercing rhs of formula to factor

## Warning in cohen.d.formula(TransformedCOEFFICIENT ~ MIGRATORY, data =
## all_provinces_mig): Cohercing rhs of formula to factor

pwr_t_tests_mig

## [1] 0.4620189 0.5431285 0.9991666 0.7756358 1.0000000

three_provinces_mig$COEFFICIENT=c(
    Ankara_mig$ANATOLIAN,
    Mugla_mig$ANATOLIAN,
    Hatay_mig$LEVANTINE)

all_provinces_mig$COEFFICIENT=c(
    Kirklareli_mig$THRACIAN,
    Edirne_mig$THRACIAN,
    Duzce_mig$ANATOLIAN,
    Eskisehir_mig$ANATOLIAN,
    Ankara_mig$ANATOLIAN,
    Mugla_mig$ANATOLIAN,
    Ardahan_mig$CAUCASIAN,
    Artvin_mig$CAUCASIAN,
    Bitlis_mig$LEVANTINE,
    Hatay_mig$LEVANTINE)

all_provinces_mig_merged = cbind(all_provinces_mig,
                                 MERGED = paste("All", all_provinces_mig$MIGR
ATORY))

all_provinces_mig$MERGED = paste(all_provinces_mig$POPULATION, all_provinces_
mig$MIGRATORY)

three_provinces_mig$MERGED = paste("Combine", three_provinces_mig$MIGRATORY)
```

```
est_fac_mig = rbind(all_provinces_mig, three_provinces_mig, all_provinces_mig
_merged)

plot(dabest(est_fac_mig, MERGED, COEFFICIENT,
         idx = list(c("Ankara 0", "Ankara 1"),
                    c("Mugla 0", "Mugla 1"),
                    c("Hatay 0", "Hatay 1"),
                    c("Combine 0", "Combine 1"),
                    c("All 0", "All 1")),
         paired = FALSE),
     theme = ggplot2::theme_gray(),
     palette = rep(c("yellow", "yellow", "violet", "coral4", "firebrick"), ea
ch= 2),
     rawplot.ylim = c(0.00, 1.10),
     effsize.ylim = c(-0.80, 0.60),
     rawplot.markersize = 4,
     rawplot.groupwidth = 0.2,
     rawplot.ylabel = "Membership Coefficient"
)
```

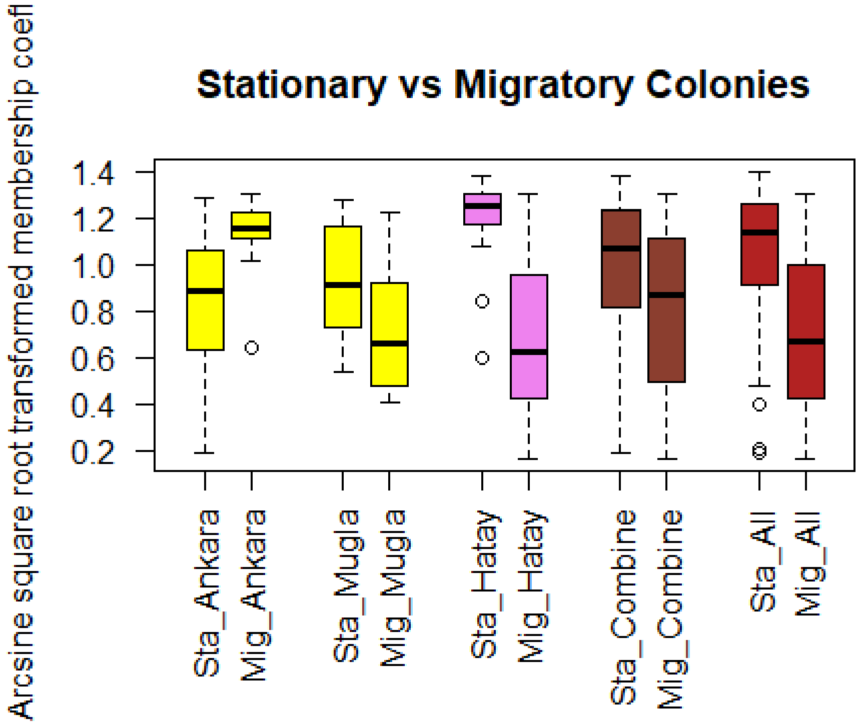


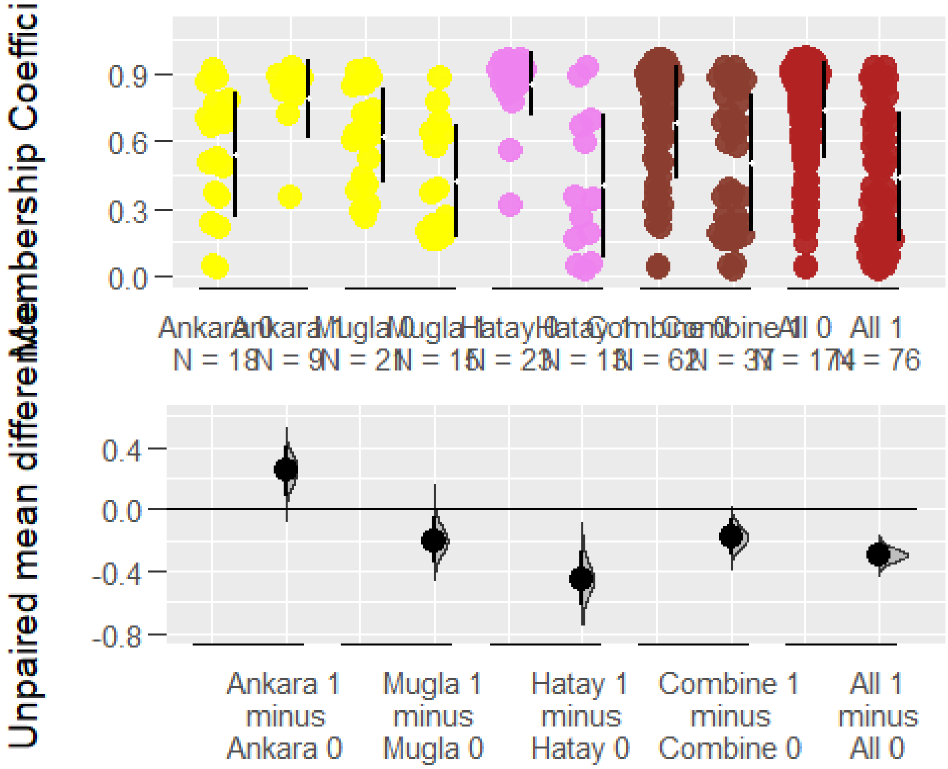


#### Isolated regions

```
Ankara_iso = subset(memberships174, memberships174$POPULATION=="Ankara")
Mugla_iso = subset(memberships174, memberships174$POPULATION=="Mugla")
Hatay_iso = subset(memberships174, memberships174$POPULATION=="Hatay")
```

```
Kirklareli_iso = subset(memberships174, memberships174$POPULATION=="Kirklarel
i")
Edirne_iso = subset(memberships174, memberships174$POPULATION=="Edirne+")
Duzce_iso = subset(memberships174, memberships174$POPULATION=="Duzce+")
Eskisehir_iso = subset(memberships174, memberships174$POPULATION=="Eskisehir+
")
Ardahan_iso = subset(memberships174, memberships174$POPULATION=="Ardahan")
Artvin_iso = subset(memberships174, memberships174$POPULATION=="Artvin")

isolated_provinces = cbind(
  rbind(
    Kirklareli_iso,
    Ardahan_iso,
    Artvin_iso),
  TransformedCOEFFICIENT=c(
    Kirklareli_iso$TransformedTHRACIAN,
    Ardahan_iso$TransformedCAUCASIAN,
    Artvin_iso$TransformedCAUCASIAN)
)

not_isolated_provinces = cbind(
  rbind(
    Edirne_iso,
    Duzce_iso,
    Eskisehir_iso,
    Ankara_iso,
    Mugla_iso,
    Hatay_iso),
  TransformedCOEFFICIENT=c(
    Edirne_iso$TransformedTHRACIAN,
    Duzce_iso$TransformedANATOLIAN,
    Eskisehir_iso$TransformedANATOLIAN,
    Ankara_iso$TransformedANATOLIAN,
    Mugla_iso$TransformedANATOLIAN,
    Hatay_iso$TransformedLEVANTINE)
)

all_provinces_iso = cbind(
  rbind(
    Kirklareli_iso,
    Edirne_iso,
    Duzce_iso,
    Eskisehir_iso,
    Ankara_iso,
    Mugla_iso,
    Ardahan_iso,
    Artvin_iso,
    Hatay_iso),
  TransformedCOEFFICIENT=c(
    Kirklareli_iso$TransformedTHRACIAN,
```

```
    Edirne_iso$TransformedTHRACIAN,
    Duzce_iso$TransformedANATOLIAN,
    Eskisehir_iso$TransformedANATOLIAN,
    Ankara_iso$TransformedANATOLIAN,
    Mugla_iso$TransformedANATOLIAN,
    Ardahan_iso$TransformedCAUCASIAN,
    Artvin_iso$TransformedCAUCASIAN,
    Hatay_iso$TransformedLEVANTINE)
)

normalities_174 = apply(memberships174[4:11], 2, function(x) {shapiro.test(x)
$p.value})
p.adjust(normalities_174, method="BH")

##            ANATOLIAN             THRACIAN            CAUCASIAN
##         4.193447e-15         1.691329e-18         3.297198e-17
##            LEVANTINE TransformedANATOLIAN  TransformedTHRACIAN
##         3.281414e-20         3.292611e-13         3.297198e-17
## TransformedCAUCASIAN TransformedLEVANTINE
##         3.349581e-15         1.691329e-18

par(mfrow=c(1,2))
boxplot(all_provinces_iso$TransformedCOEFFICIENT ~ ordered(all_provinces_iso$
POPULATION, levels=rev(c("Kirklareli", "Edirne+", "Duzce+", "Eskisehir+", "Mu
gla", "Ankara", "Ardahan", "Artvin", "Hatay"))), data = all_provinces_iso,
        col= rep(rev(c("orangered", "yellow", "blue", "violet")), times= rev(
c(2,4,2,1))),
        horizontal= TRUE,
        las=1,
        xlab="Transformed membership coefficients",
        ylab="",
        pars=list(par(mar=c(8,6,4,2))),
        at =rev(c(12,11, 9,8,7,6, 4,3, 1))
)
boxplot(
  not_isolated_provinces$TransformedCOEFFICIENT,
  isolated_provinces$TransformedCOEFFICIENT,
  col=c("forestgreen","darkorchid3"),
  horizontal= TRUE,
  las=1,
  xlab="Transformed membership coefficients",
  ylab="",
  pars=list(par(mar=c(8,0,4,2)))
)
legend("left",
       inset= 0.01,
       c("isolated", "not_isolated"),
       fill=c("darkorchid3", "forestgreen"),
       cex=0.7,
       text.font=2,
```

```
       box.lty=0,
       horiz=FALSE)
```

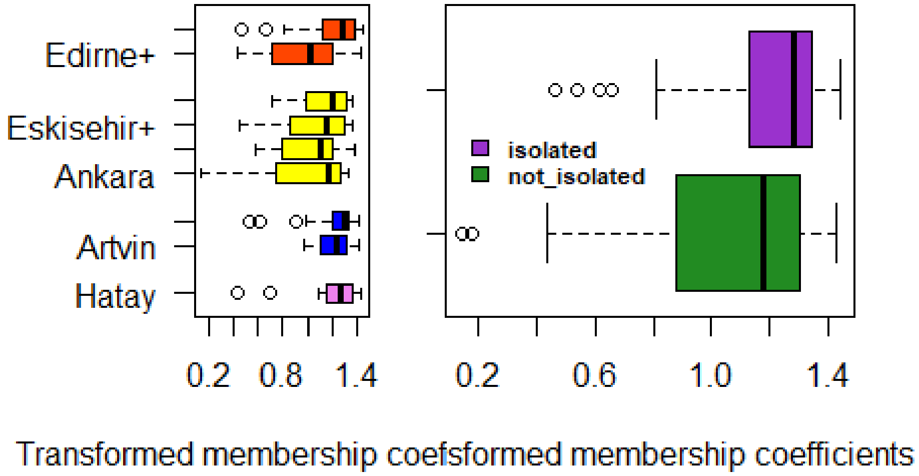


```
par(mfrow=c(1,1))

mean_iso = apply(cbind.na(
  isolated_provinces$TransformedCOEFFICIENT,
  not_isolated_provinces$TransformedCOEFFICIENT),
  2, function(x) {mean(x, na.rm=TRUE)})
mean_iso

## [1] 1.210052 1.083340

cohen.d(isolated_provinces$TransformedCOEFFICIENT, not_isolated_provinces$Tra
nsformedCOEFFICIENT)$estimate

## [1] 0.4897751

wilcox.test(isolated_provinces$TransformedCOEFFICIENT, not_isolated_provinces
$TransformedCOEFFICIENT)$p.value

## [1] 0.002100692

var.test(isolated_provinces$TransformedCOEFFICIENT, not_isolated_provinces$Tr
ansformedCOEFFICIENT)$p.value

## [1] 0.002916549
```

```r
t.test(isolated_provinces$TransformedCOEFFICIENT, not_isolated_provinces$Tran
sformedCOEFFICIENT, var.equal = FALSE)$p.value

## [1] 0.001130164

pwr.t2n.test(
  n1= length(isolated_provinces$TransformedCOEFFICIENT),
  n2= length(not_isolated_provinces$TransformedCOEFFICIENT),
  d= cohen.d(isolated_provinces$TransformedCOEFFICIENT, not_isolated_province
s$TransformedCOEFFICIENT)$estimate,
  sig.level= 0.05)$power

## [1] 0.8922584

all_provinces_iso$COEFFICIENT=c(
    Kirklareli_iso$THRACIAN,
    Edirne_iso$THRACIAN,
    Duzce_iso$ANATOLIAN,
    Eskisehir_iso$ANATOLIAN,
    Ankara_iso$ANATOLIAN,
    Mugla_iso$ANATOLIAN,
    Ardahan_iso$CAUCASIAN,
    Artvin_iso$CAUCASIAN,
    Hatay_iso$LEVANTINE)

plot(dabest(all_provinces_iso, POPULATION, COEFFICIENT,
         idx = c("Kirklareli", "Edirne+", "Duzce+", "Eskisehir+", "Mugla", "A
nkara", "Ardahan", "Artvin", "Hatay"),
         paired = FALSE),
     theme = ggplot2::theme_gray(),
     palette = rep(c("orangered", "yellow", "blue", "violet"), times= c(2,4,2
,1)),
     rawplot.ylim = c(0.00, 1.10),
     effsize.ylim = c(-0.70, 0.30),
     rawplot.markersize = 4,
     rawplot.groupwidth = 0.2,
     rawplot.ylabel = "Membership Coefficient",
     axes.title.fontsize = 12
)
```

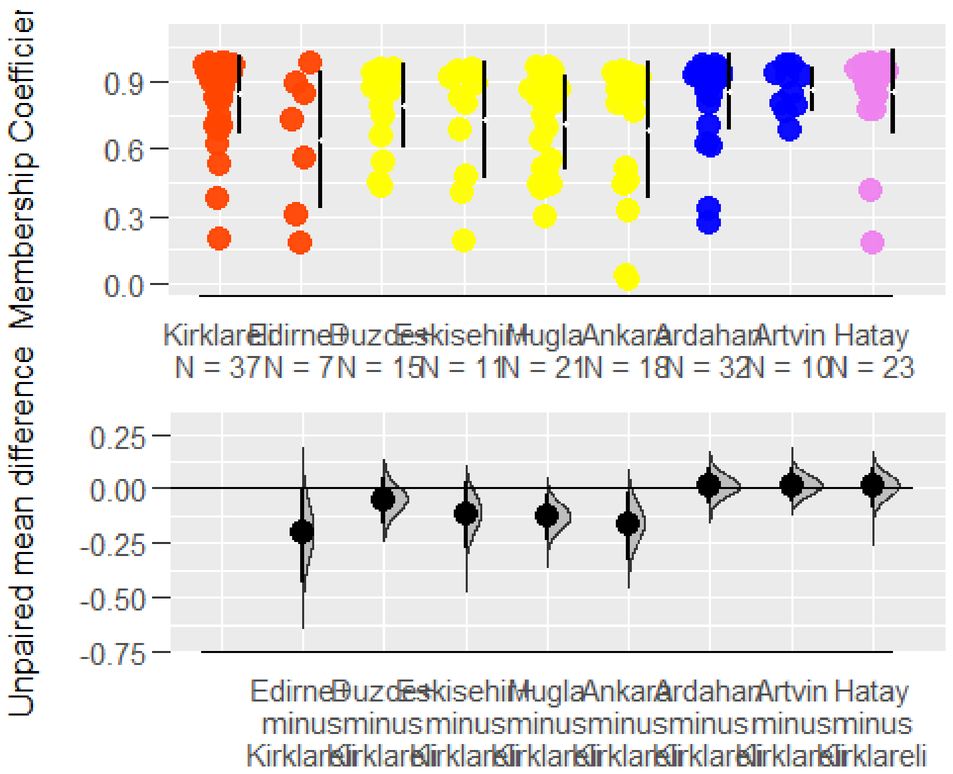


```
isolated_provinces$COEFFICIENT=c(
    Kirklareli_iso$THRACIAN,
    Ardahan_iso$CAUCASIAN,
    Artvin_iso$CAUCASIAN)

not_isolated_provinces$COEFFICIENT=c(
    Edirne_iso$THRACIAN,
    Duzce_iso$ANATOLIAN,
    Eskisehir_iso$ANATOLIAN,
    Ankara_iso$ANATOLIAN,
    Mugla_iso$ANATOLIAN,
    Hatay_iso$LEVANTINE)

est_fac_iso = rbind(
  cbind.data.frame(COEFFICIENT = isolated_provinces$COEFFICIENT, Isolated = "
1"),
  cbind.data.frame(COEFFICIENT = not_isolated_provinces$COEFFICIENT, Isolated
= "0")
)

plot(dabest(est_fac_iso, Isolated, COEFFICIENT,
         idx = c("0", "1"),
         paired = FALSE),
     theme = ggplot2::theme_gray(),
     palette = c("forestgreen", "darkorchid3"),
     rawplot.ylim = c(0.00, 1.10),
```

```
     rawplot.markersize = 4,
     rawplot.groupwidth = 0.2,
     rawplot.ylabel = "Membership Coefficient"
)
```

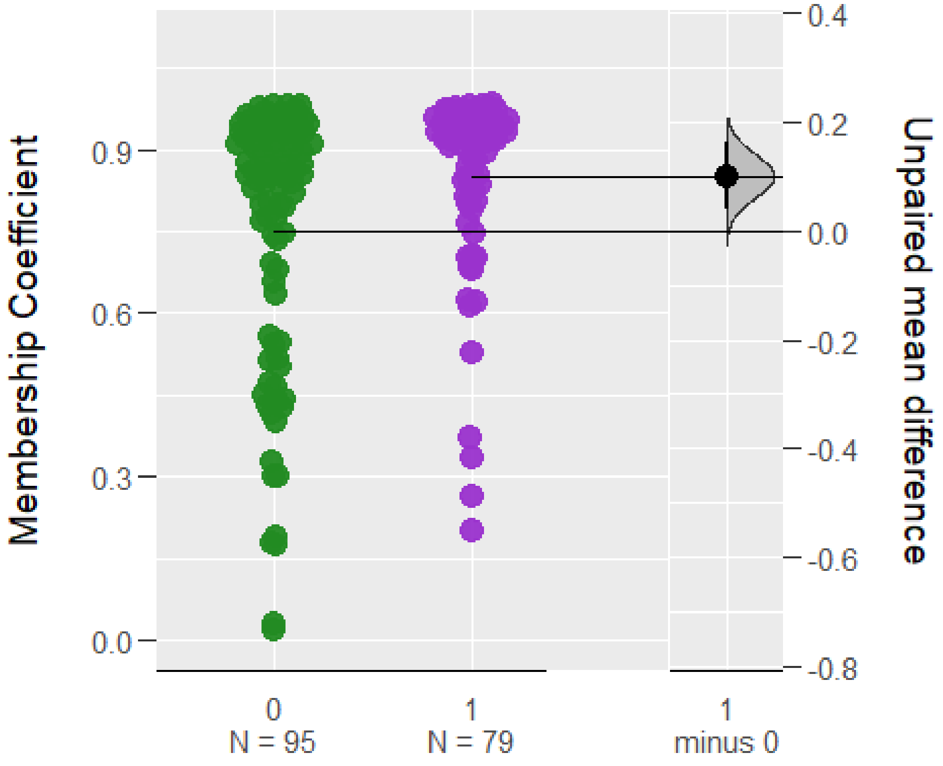


#### Queen and colony trade

```
Thracian_mis = cbind(
  rbind(
    Duzce_iso,
    Eskisehir_iso,
    Ankara_iso,
    Mugla_iso,
    Ardahan_iso,
    Artvin_iso,
    Hatay_iso),
  TransformedCOEFFICIENT=c(
    Duzce_iso$TransformedTHRACIAN,
    Eskisehir_iso$TransformedTHRACIAN,
    Ankara_iso$TransformedTHRACIAN,
    Mugla_iso$TransformedTHRACIAN,
    Ardahan_iso$TransformedTHRACIAN,
    Artvin_iso$TransformedTHRACIAN,
    Hatay_iso$TransformedTHRACIAN)
)

Anatolian_mis = cbind(
```

```
  rbind(
    Kirklareli_iso,
    Edirne_iso,
    Ardahan_iso,
    Artvin_iso,
    Hatay_iso),
  TransformedCOEFFICIENT=c(
    Kirklareli_iso$TransformedANATOLIAN,
    Edirne_iso$TransformedANATOLIAN,
    Ardahan_iso$TransformedANATOLIAN,
    Artvin_iso$TransformedANATOLIAN,
    Hatay_iso$TransformedANATOLIAN)
)

Caucasian_mis = cbind(
  rbind(
    Kirklareli_iso,
    Edirne_iso,
    Duzce_iso,
    Eskisehir_iso,
    Ankara_iso,
    Mugla_iso,
    Hatay_iso),
  TransformedCOEFFICIENT=c(
    Kirklareli_iso$TransformedCAUCASIAN,
    Edirne_iso$TransformedCAUCASIAN,
    Duzce_iso$TransformedCAUCASIAN,
    Eskisehir_iso$TransformedCAUCASIAN,
    Ankara_iso$TransformedCAUCASIAN,
    Mugla_iso$TransformedCAUCASIAN,
    Hatay_iso$TransformedCAUCASIAN)
)

Levantine_mis = cbind(
  rbind(
    Kirklareli_iso,
    Edirne_iso,
    Duzce_iso,
    Eskisehir_iso,
    Ankara_iso,
    Mugla_iso,
    Ardahan_iso,
    Artvin_iso),
  TransformedCOEFFICIENT=c(
    Kirklareli_iso$TransformedLEVANTINE,
    Edirne_iso$TransformedLEVANTINE,
    Duzce_iso$TransformedLEVANTINE,
    Eskisehir_iso$TransformedLEVANTINE,
    Ankara_iso$TransformedLEVANTINE,
    Mugla_iso$TransformedLEVANTINE,
```

```
    Ardahan_iso$TransformedLEVANTINE,
    Artvin_iso$TransformedLEVANTINE)
)

boxplot(list(
  Thracian = Thracian_mis$TransformedCOEFFICIENT,
  Anatolian = Anatolian_mis$TransformedCOEFFICIENT,
  Caucasian = Caucasian_mis$TransformedCOEFFICIENT,
  Levantine = Levantine_mis$TransformedCOEFFICIENT),
  col=c("orangered","yellow","blue","violet"),
  las=1,
  xlab="",
  ylab="Arcsine square root transformed membership coefficients",
  pars=list(par(mar=c(4,6,4,2))),
  main="Misassignments to Clusters"
)
```

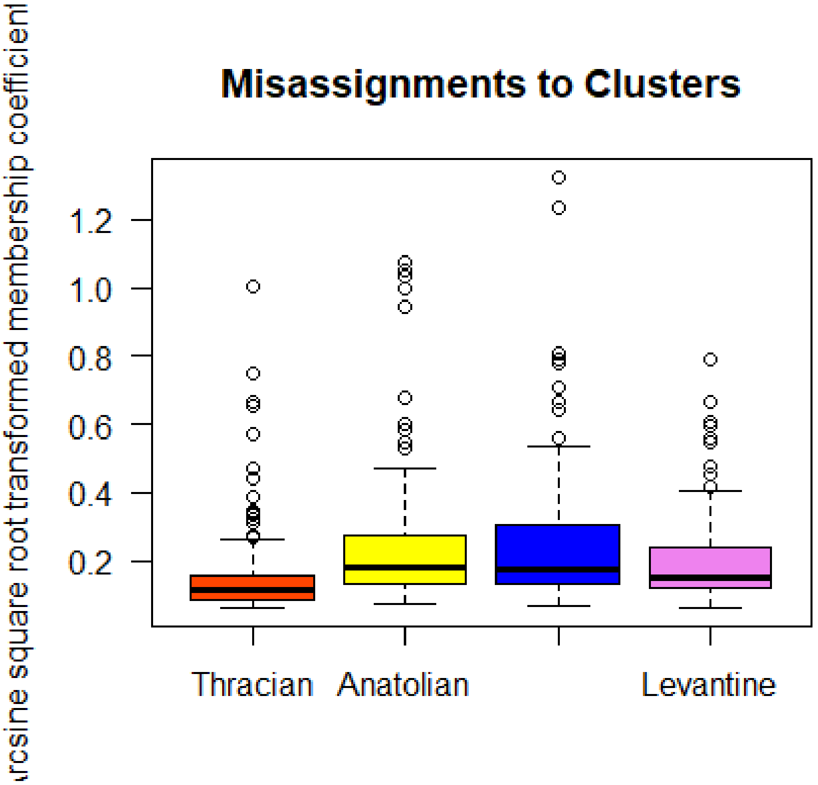


```
mean_mis = apply(cbind.na(
  Thracian_mis$TransformedCOEFFICIENT,
  Anatolian_mis$TransformedCOEFFICIENT,
  Caucasian_mis$TransformedCOEFFICIENT,
  Levantine_mis$TransformedCOEFFICIENT),
  2, function(x) {mean(x, na.rm=TRUE)})
mean_mis

## [1] 0.1599180 0.2546807 0.2576699 0.1976263
```

```
effect_sizes_mis = c(
  cohen.d(Thracian_mis$TransformedCOEFFICIENT, Anatolian_mis$TransformedCOEFF
ICIENT)$estimate,
  cohen.d(Thracian_mis$TransformedCOEFFICIENT, Caucasian_mis$TransformedCOEFF
ICIENT)$estimate,
  cohen.d(Anatolian_mis$TransformedCOEFFICIENT, Caucasian_mis$TransformedCOEF
FICIENT)$estimate,
  cohen.d(Thracian_mis$TransformedCOEFFICIENT, Levantine_mis$TransformedCOEFF
ICIENT)$estimate,
  cohen.d(Anatolian_mis$TransformedCOEFFICIENT, Levantine_mis$TransformedCOEF
FICIENT)$estimate,
  cohen.d(Caucasian_mis$TransformedCOEFFICIENT, Levantine_mis$TransformedCOEF
FICIENT)$estimate
)
effect_sizes_mis

## [1] -0.53178083 -0.53846589 -0.01395682 -0.27940263  0.33515954  0.3448442
8

kruskal.test(list(
  Thracian_mis$TransformedCOEFFICIENT,
  Anatolian_mis$TransformedCOEFFICIENT,
  Caucasian_mis$TransformedCOEFFICIENT,
  Levantine_mis$TransformedCOEFFICIENT),
  method="bh")$p.value

## [1] 8.161421e-12

dunn.test(list(
  Thracian_mis$TransformedCOEFFICIENT,
  Anatolian_mis$TransformedCOEFFICIENT,
  Caucasian_mis$TransformedCOEFFICIENT,
  Levantine_mis$TransformedCOEFFICIENT),
  method="bh", table=FALSE, list=TRUE, altp=TRUE)$altP.adjust

##   Kruskal-Wallis rank sum test
##
## data: x and group
## Kruskal-Wallis chi-squared = 54.6482, df = 3, p-value = 0
##
##
##                            Comparison of x by group
##                              (Benjamini-Hochberg)
##
## List of pairwise comparisons: Z statistic (adjusted p-value)
## ---------------------------
## 1 - 2 : -6.185241 (0.0000)*
## 1 - 3 : -6.517906 (0.0000)*
## 2 - 3 : -0.016162 (0.9871)
## 1 - 4 : -4.392869 (0.0000)*
## 2 - 4 :  2.209515 (0.0326)*
```

```
## 3 - 4 :  2.348135 (0.0283)*
##
## alpha = 0.05
## Reject Ho if p <= alpha
```

```
## [1] 1.860224e-09 4.277724e-10 9.871048e-01 2.237283e-05 3.256659e-02
## [6] 2.830147e-02
```

```
anova_fac_mis = rbind(
  cbind.data.frame(TransformedCOEFFICIENT = Thracian_mis$TransformedCOEFFICIE
NT, Origin = gsub("_.*$", "", "Thracian_mis$TransformedCOEFFICIENT")),
  cbind.data.frame(TransformedCOEFFICIENT = Anatolian_mis$TransformedCOEFFICI
ENT, Origin = gsub("_.*$", "", "Anatolian_mis$TransformedCOEFFICIENT")),
  cbind.data.frame(TransformedCOEFFICIENT = Caucasian_mis$TransformedCOEFFICI
ENT, Origin = gsub("_.*$", "", "Caucasian_mis$TransformedCOEFFICIENT")),
  cbind.data.frame(TransformedCOEFFICIENT = Levantine_mis$TransformedCOEFFICI
ENT, Origin = gsub("_.*$", "", "Levantine_mis$TransformedCOEFFICIENT"))
)
```

```
summary(aov(TransformedCOEFFICIENT ~ Origin, anova_fac_mis))[[1]][["Pr(>F)"]]
[[1]]
```

```
## [1] 6.410609e-06
```

```
TukeyHSD(aov(TransformedCOEFFICIENT ~ Origin, anova_fac_mis))$Origin[,4]
```

```
## Caucasian-Anatolian Levantine-Anatolian  Thracian-Anatolian Levantine-Cauc
asian
##        9.991923e-01        4.975158e-02        2.308165e-04        2.25857
2e-02
##  Thracian-Caucasian  Thracian-Levantine
##        5.070592e-05        2.789071e-01
```

```
plot(jitter(Thracian_mis$TransformedCOEFFICIENT,amount=0.05), pch=21, bg="ora
ngered", cex=1.5, col="black", xlim=c(0, 151), ylim=c(0, 1.5), las=1,
  xlab="",
  ylab="Arcsine square root transformed membership coefficients",
  pars=list(par(mar=c(4,6,4,2))),
  main="Individual Misassignments")
```

```
## Warning in plot.window(...): "pars" bir grafiksel parametre değil
```

```
## Warning in plot.xy(xy, type, ...): "pars" bir grafiksel parametre değil
```

```
## Warning in axis(side = side, at = at, labels = labels, ...): "pars" bir
## grafiksel parametre değil
```

```
## Warning in axis(side = side, at = at, labels = labels, ...): "pars" bir
## grafiksel parametre değil
```

```
## Warning in box(...): "pars" bir grafiksel parametre değil
```

```
## Warning in title(...): "pars" bir grafiksel parametre değil
```

```
points(jitter(Anatolian_mis$TransformedCOEFFICIENT,amount=0.05), pch=21, bg="
yellow", cex=1.5, col="black")
points(jitter(Caucasian_mis$TransformedCOEFFICIENT,amount=0.05), pch=21, bg="
blue", cex=1.5, col="black")
points(jitter(Levantine_mis$TransformedCOEFFICIENT,amount=0.05), pch=21, bg="
violet", cex=1.5, col="black")
abline(h=0.5, col="black",lwd=4)
```

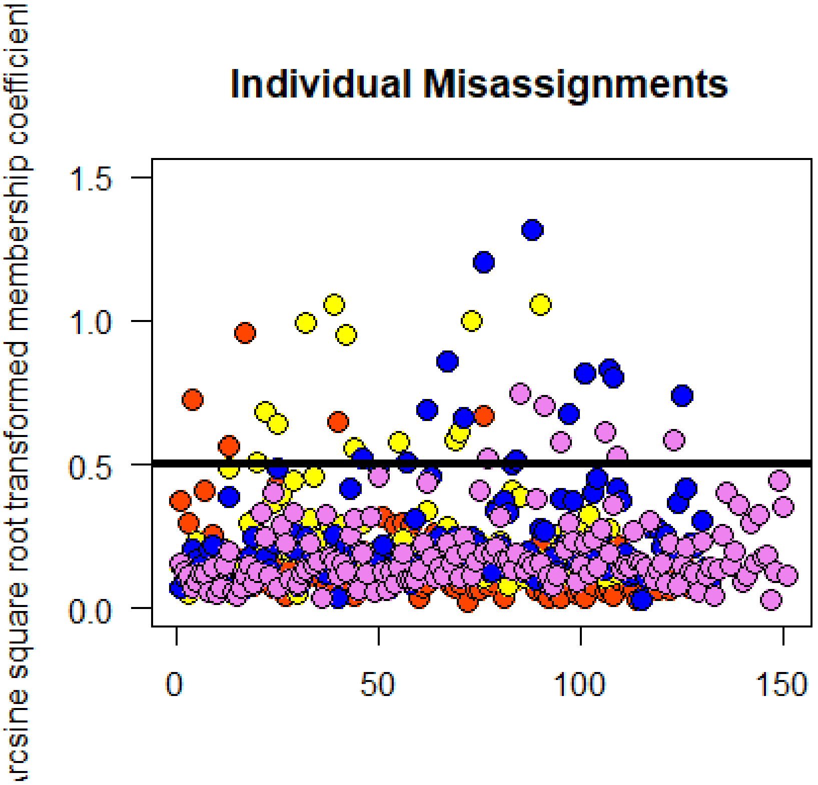


```
large_mis = sum(cbind.na(
  Thracian_mis$TransformedCOEFFICIENT,
  Anatolian_mis$TransformedCOEFFICIENT,
  Caucasian_mis$TransformedCOEFFICIENT,
  Levantine_mis$TransformedCOEFFICIENT)>=0.5, na.rm=TRUE)/
            sum(cbind.na(
  Thracian_mis$TransformedCOEFFICIENT,
  Anatolian_mis$TransformedCOEFFICIENT,
  Caucasian_mis$TransformedCOEFFICIENT,
  Levantine_mis$TransformedCOEFFICIENT)>0, na.rm=TRUE)
large_mis
```

```
## [1] 0.07471264
```

```
par(mfrow=c(2,2))
boxplot(TransformedTHRACIAN ~ ordered(Thracian_mis$POPULATION, levels=c("Kirk
lareli", "Edirne+", "Duzce+", "Eskisehir+", "Mugla", "Ankara", "Ardahan", "Ar
tvin", "Hatay")),
```

```
        data=Thracian_mis,
        col=c("orangered"),
        las=1,
        xlab="",
        ylab="",
        pars=list(par(mar=c(4,4,4,0))),
        main="Misassignments to Thracian Cluster",
        ylim=c(0, 1.4),
        names = c("Kirklareli", "Edirne+", "Duzce+", "Eskisehir+", "Mugla", "
Ankara", "Ardahan", "Artvin", "Hatay")
)
boxplot(TransformedANATOLIAN ~ ordered(Anatolian_mis$POPULATION, levels=c("Ki
rklareli", "Edirne+", "Duzce+", "Eskisehir+", "Mugla", "Ankara", "Ardahan", "
Artvin", "Hatay")),
        data=Anatolian_mis,
        col=c("yellow"),
        las=1,
        xlab="",
        ylab="",
        pars=list(par(mar=c(4,2.5,4,1.5))),
        main="Misassignments to Anatolian Cluster",
        ylim=c(0, 1.4),
        names = c("Kirklareli", "Edirne+", "Duzce+", "Eskisehir+", "Mugla", "
Ankara", "Ardahan", "Artvin", "Hatay")
)
boxplot(TransformedCAUCASIAN ~ ordered(Caucasian_mis$POPULATION, levels=c("Ki
rklareli", "Edirne+", "Duzce+", "Eskisehir+", "Mugla", "Ankara", "Ardahan", "
Artvin", "Hatay")),
        data=Caucasian_mis,
        col=c("blue"),
        las=1,
        xlab="",
        ylab="",
        pars=list(par(mar=c(4,4,4,0))),
        main="Misassignments to Caucasian Cluster",
        ylim=c(0, 1.4),
        names = c("Kirklareli", "Edirne+", "Duzce+", "Eskisehir+", "Mugla", "
Ankara", "Ardahan", "Artvin", "Hatay")
)
boxplot(TransformedLEVANTINE ~ ordered(Levantine_mis$POPULATION, levels=c("Ki
rklareli", "Edirne+", "Duzce+", "Eskisehir+", "Mugla", "Ankara", "Ardahan", "
Artvin", "Hatay")),
        data=Levantine_mis,
        col=c("violet"),
        las=1,
        xlab="",
        ylab="",
        pars=list(par(mar=c(4,2.5,4,1.5))),
        main="Misassignments to Levantine Cluster",
        ylim=c(0, 1.4),
```

```r
        names = c("Kirklareli", "Edirne+", "Duzce+", "Eskisehir+", "Mugla", "
Ankara", "Ardahan", "Artvin", "Hatay")
)
par(mfrow=c(1,1))
mtext("Arcsine square root transformed membership coefficients", side = 2, li
ne = 1.5)
```

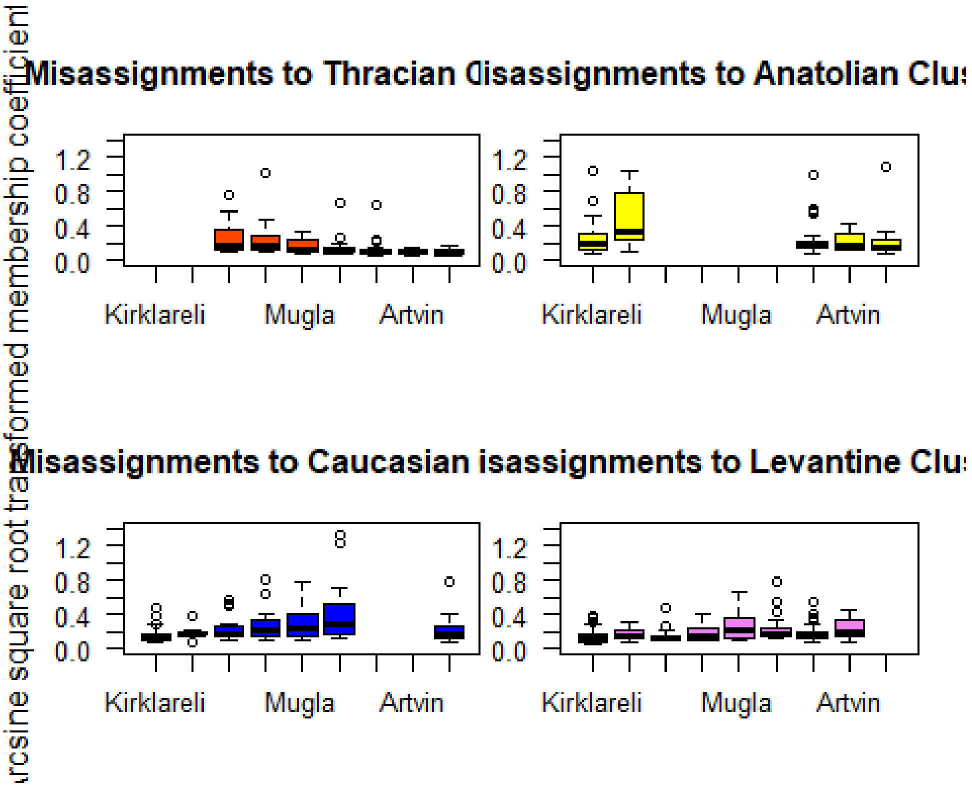


```r
dunn.test(Thracian_mis$TransformedTHRACIAN, Thracian_mis$POPULATION, method="
bh", table=FALSE, list=TRUE, altp=TRUE)$altP.adjust

##   Kruskal-Wallis rank sum test
##
## data: x and group
## Kruskal-Wallis chi-squared = 33.7717, df = 6, p-value = 0
##
##
##                             Comparison of x by group
##                               (Benjamini-Hochberg)
##
## List of pairwise comparisons: Z statistic (adjusted p-value)
## ------------------------------------------
## Ankara - Ardahan     :  0.733480 (0.5405)
## Ankara - Artvin      :  1.187973 (0.3288)
## Ardahan - Artvin     :  0.696798 (0.5371)
## Ankara - Duzce+      : -2.995549 (0.0096)*
## Ardahan - Duzce+     : -4.037362 (0.0006)*
## Artvin - Duzce+      : -3.712926 (0.0014)*
```

```
## Ankara - Eskisehir+  : -2.180257 (0.0682)
## Ardahan - Eskisehir+ : -3.005623 (0.0111)*
## Artvin - Eskisehir+  : -2.982029 (0.0086)*
## Duzce+ - Eskisehir+  :  0.536211 (0.6214)
## Ankara - Hatay       :  1.346935 (0.2875)
## Ardahan - Hatay      :  0.760054 (0.5525)
## Artvin - Hatay       : -0.117921 (0.9061)
## Duzce+ - Hatay       :  4.432701 (0.0002)*
## Eskisehir+ - Hatay   :  3.432389 (0.0031)*
## Ankara - Mugla       : -1.138340 (0.3347)
## Ardahan - Mugla      : -2.071484 (0.0731)
## Artvin - Mugla       : -2.171162 (0.0628)
## Duzce+ - Mugla       :  2.016226 (0.0766)
## Eskisehir+ - Mugla   :  1.259439 (0.3118)
## Hatay - Mugla        : -2.615835 (0.0234)*
##
## alpha = 0.05
## Reject Ho if p <= alpha
```

```
##  [1] 0.5404762394 0.3287816257 0.5370795810 0.0095882641 0.0005675828
##  [6] 0.0014341333 0.0682229193 0.0111315412 0.0085903221 0.6214032526
## [11] 0.2875399886 0.5524509169 0.9061294848 0.0001954254 0.0031410113
## [16] 0.3346588667 0.0731440909 0.0628297065 0.0766085639 0.3118073401
## [21] 0.0233649965
```

```
dunn.test(Anatolian_mis$TransformedANATOLIAN, Anatolian_mis$POPULATION, metho
d="bh", table=FALSE, list=TRUE, altp=TRUE)$altP.adjust
```

```
##   Kruskal-Wallis rank sum test
##
## data: x and group
## Kruskal-Wallis chi-squared = 6.2956, df = 4, p-value = 0.18
##
##
##                           Comparison of x by group
##                             (Benjamini-Hochberg)
##
## List of pairwise comparisons: Z statistic (adjusted p-value)
## ----------------------------------------
## Ardahan - Artvin     : -0.319555 (0.8326)
## Ardahan - Edirne+    : -2.152900 (0.1566)
## Artvin - Edirne+     : -1.587954 (0.2807)
## Ardahan - Hatay      :  0.608762 (0.7753)
## Artvin - Hatay       :  0.744971 (0.7605)
## Edirne+ - Hatay      :  2.466576 (0.1364)
## Ardahan - Kirklareli : -0.409257 (0.8529)
## Artvin - Kirklareli  :  0.047621 (0.9620)
## Edirne+ - Kirklareli :  1.939794 (0.1747)
## Hatay - Kirklareli   : -0.998807 (0.6358)
##
```

```
## alpha = 0.05
## Reject Ho if p <= alpha

##  [1] 0.8325614 0.1566324 0.2807415 0.7752596 0.7604815 0.1364116 0.8529382
##  [8] 0.9620182 0.1746823 0.6357759
```

```r
dunn.test(Caucasian_mis$TransformedCAUCASIAN, Caucasian_mis$POPULATION, metho
d="bh", table=FALSE, list=TRUE, altp=TRUE)$altP.adjust
```

```
##   Kruskal-Wallis rank sum test
##
## data: x and group
## Kruskal-Wallis chi-squared = 24.4867, df = 6, p-value = 0
##
##
##                            Comparison of x by group
##                              (Benjamini-Hochberg)
##
## List of pairwise comparisons: Z statistic (adjusted p-value)
## ---------------------------------------------
## Ankara - Duzce+          :  1.503982 (0.3480)
## Ankara - Edirne+         :  1.462703 (0.3349)
## Duzce+ - Edirne+         :  0.274705 (0.8227)
## Ankara - Eskisehir+      :  0.941227 (0.5599)
## Duzce+ - Eskisehir+      : -0.417128 (0.7478)
## Edirne+ - Eskisehir+     : -0.602542 (0.7177)
## Ankara - Hatay           :  2.156034 (0.1632)
## Duzce+ - Hatay           :  0.460103 (0.7973)
## Edirne+ - Hatay          :  0.062447 (0.9502)
## Eskisehir+ - Hatay       :  0.868226 (0.5779)
## Ankara - Kirklareli      :  4.116834 (0.0008)*
## Duzce+ - Kirklareli      :  2.147265 (0.1334)
## Edirne+ - Kirklareli     :  1.289570 (0.4141)
## Eskisehir+ - Kirklareli  :  2.396051 (0.1160)
## Hatay - Kirklareli       :  1.900234 (0.2009)
## Ankara - Mugla           :  0.417954 (0.7886)
## Duzce+ - Mugla           : -1.158209 (0.4319)
## Edirne+ - Mugla          : -1.185258 (0.4504)
## Eskisehir+ - Mugla       : -0.607113 (0.7613)
## Hatay - Mugla            : -1.803193 (0.2141)
## Kirklareli - Mugla       : -3.838787 (0.0013)*
##
## alpha = 0.05
## Reject Ho if p <= alpha

##  [1] 0.3480379226 0.3349468384 0.8227197142 0.5598734459 0.7478041560
##  [6] 0.7176924243 0.1631751049 0.7973106498 0.9502063240 0.5779059142
## [11] 0.0008066361 0.1334431588 0.4141193923 0.1160093157 0.2009084083
## [16] 0.7886436758 0.4318623501 0.4503834466 0.7612855064 0.2140734064
## [21] 0.0012982542
```

```
dunn.test(Levantine_mis$TransformedLEVANTINE, Levantine_mis$POPULATION, metho
d="bh", table=FALSE, list=TRUE, altp=TRUE)$altP.adjust
```

```
##   Kruskal-Wallis rank sum test
##
## data: x and group
## Kruskal-Wallis chi-squared = 19.2166, df = 7, p-value = 0.01
##
##
##                            Comparison of x by group
##                              (Benjamini-Hochberg)
##
## List of pairwise comparisons: Z statistic (adjusted p-value)
## ---------------------------------------------
## Ankara - Ardahan         :  1.827729 (0.2704)
## Ankara - Artvin          :  0.462194 (0.8196)
## Ardahan - Artvin         : -0.983227 (0.5361)
## Ankara - Duzce+          :  2.477045 (0.0927)
## Ardahan - Duzce+         :  1.046549 (0.5906)
## Artvin - Duzce+          :  1.674692 (0.3290)
## Ankara - Edirne+         :  1.556922 (0.3346)
## Ardahan - Edirne+        :  0.371489 (0.8287)
## Artvin - Edirne+         :  1.037359 (0.5592)
## Duzce+ - Edirne+         : -0.376797 (0.8599)
## Ankara - Eskisehir+      :  1.437719 (0.3242)
## Ardahan - Eskisehir+     :  0.033547 (0.9732)
## Artvin - Eskisehir+      :  0.842084 (0.6218)
## Duzce+ - Eskisehir+      : -0.795442 (0.6283)
## Edirne+ - Eskisehir+     : -0.296349 (0.8590)
## Ankara - Kirklareli      :  3.121411 (0.0252)*
## Ardahan - Kirklareli     :  1.485072 (0.3209)
## Artvin - Kirklareli      :  2.005322 (0.2516)
## Duzce+ - Kirklareli      :  0.101352 (0.9900)
## Edirne+ - Kirklareli     :  0.493722 (0.8287)
## Eskisehir+ - Kirklareli  :  1.009791 (0.5470)
## Ankara - Mugla           : -0.040960 (1.0000)
## Ardahan - Mugla          : -1.964335 (0.2310)
## Artvin - Mugla           : -0.508700 (0.8553)
## Duzce+ - Mugla           : -2.600529 (0.0869)
## Edirne+ - Mugla          : -1.619173 (0.3279)
## Eskisehir+ - Mugla       : -1.513679 (0.3312)
## Kirklareli - Mugla       : -3.331314 (0.0242)*
##
## alpha = 0.05
## Reject Ho if p <= alpha
```

```
##  [1] 0.27036080 0.81956219 0.53611055 0.09273248 0.59061459 0.32898092
##  [7] 0.33456913 0.82865162 0.55919450 0.85987322 0.32418351 0.97323800
## [13] 0.62181896 0.62831448 0.85899888 0.02519806 0.32089054 0.25160027
```

```
## [19] 0.98998364 0.82866946 0.54704185 1.00000000 0.23095879 0.85534693
## [25] 0.08687469 0.32794211 0.33118153 0.02420233
```

```
Thracian_mis$MISASSIGN = "Thracian"
Anatolian_mis$MISASSIGN = "Anatolian"
Caucasian_mis$MISASSIGN = "Caucasian"
Levantine_mis$MISASSIGN = "Levantine"

Thracian_mis$COEFFICIENT=c(
    Duzce_iso$THRACIAN,
    Eskisehir_iso$THRACIAN,
    Ankara_iso$THRACIAN,
    Mugla_iso$THRACIAN,
    Ardahan_iso$THRACIAN,
    Artvin_iso$THRACIAN,
    Hatay_iso$THRACIAN)

Anatolian_mis$COEFFICIENT=c(
    Kirklareli_iso$ANATOLIAN,
    Edirne_iso$ANATOLIAN,
    Ardahan_iso$ANATOLIAN,
    Artvin_iso$ANATOLIAN,
    Hatay_iso$ANATOLIAN)

Caucasian_mis$COEFFICIENT=c(
    Kirklareli_iso$CAUCASIAN,
    Edirne_iso$CAUCASIAN,
    Duzce_iso$CAUCASIAN,
    Eskisehir_iso$CAUCASIAN,
    Ankara_iso$CAUCASIAN,
    Mugla_iso$CAUCASIAN,
    Hatay_iso$CAUCASIAN)

Levantine_mis$COEFFICIENT=c(
    Kirklareli_iso$LEVANTINE,
    Edirne_iso$LEVANTINE,
    Duzce_iso$LEVANTINE,
    Eskisehir_iso$LEVANTINE,
    Ankara_iso$LEVANTINE,
    Mugla_iso$LEVANTINE,
    Ardahan_iso$LEVANTINE,
    Artvin_iso$LEVANTINE)

est_fac_iso_mis = rbind(Thracian_mis, Anatolian_mis, Caucasian_mis, Levantine
_mis)

plot(dabest(est_fac_iso_mis, MISASSIGN, COEFFICIENT,
         idx = c("Thracian", "Anatolian", "Caucasian", "Levantine"),
         paired = FALSE),
     theme = ggplot2::theme_gray(),
```

```
    palette = c("orangered", "yellow", "blue", "violet"),
    rawplot.ylim = c(0.00, 1.10),
    effsize.ylim = c(-0.20, 0.20),
    rawplot.markersize = 4,
    rawplot.groupwidth = 0.2,
    rawplot.ylabel = "Membership Coefficient"
)
```

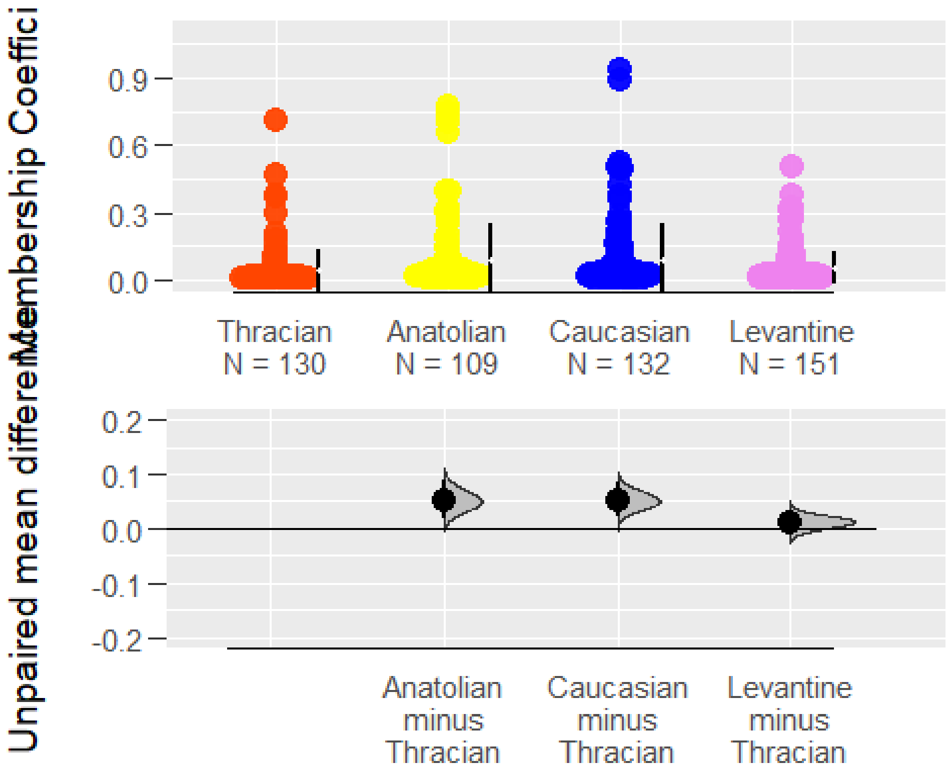


```
plot(dabest(all_provinces_iso, POPULATION, THRACIAN,
         idx = c("Kirklareli", "Edirne+", "Duzce+", "Eskisehir+", "Mugla", "A
nkara", "Ardahan", "Artvin", "Hatay"),
         paired = FALSE),
    theme = ggplot2::theme_gray(),
    palette = rep(c("orangered"), times= 9),
    rawplot.ylim = c(0.00, 1.10),
    effsize.ylim = c(-1.00, 0.40),
    rawplot.markersize = 4,
    rawplot.groupwidth = 0.2,
    rawplot.ylabel = "Membership Coefficient",
    axes.title.fontsize = 12
)
```

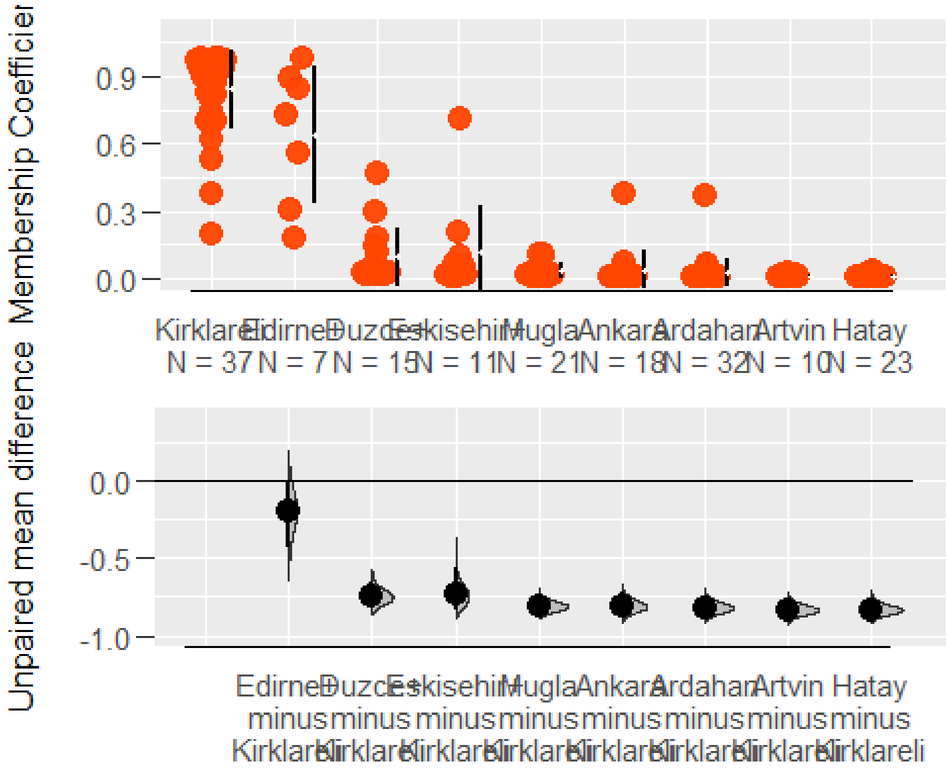


```
plot(dabest(all_provinces_iso, POPULATION, ANATOLIAN,
         idx = c("Duzce+", "Eskisehir+", "Mugla", "Ankara", "Kirklareli", "Ed
irne+", "Ardahan", "Artvin", "Hatay"),
         paired = FALSE),
     theme = ggplot2::theme_gray(),
     palette = rep(c("yellow"), times= 9),
     rawplot.ylim = c(0.00, 1.10),
     effsize.ylim = c(-1.00, 0.40),
     rawplot.markersize = 4,
     rawplot.groupwidth = 0.2,
     rawplot.ylabel = "Membership Coefficient",
     axes.title.fontsize = 12
)
```

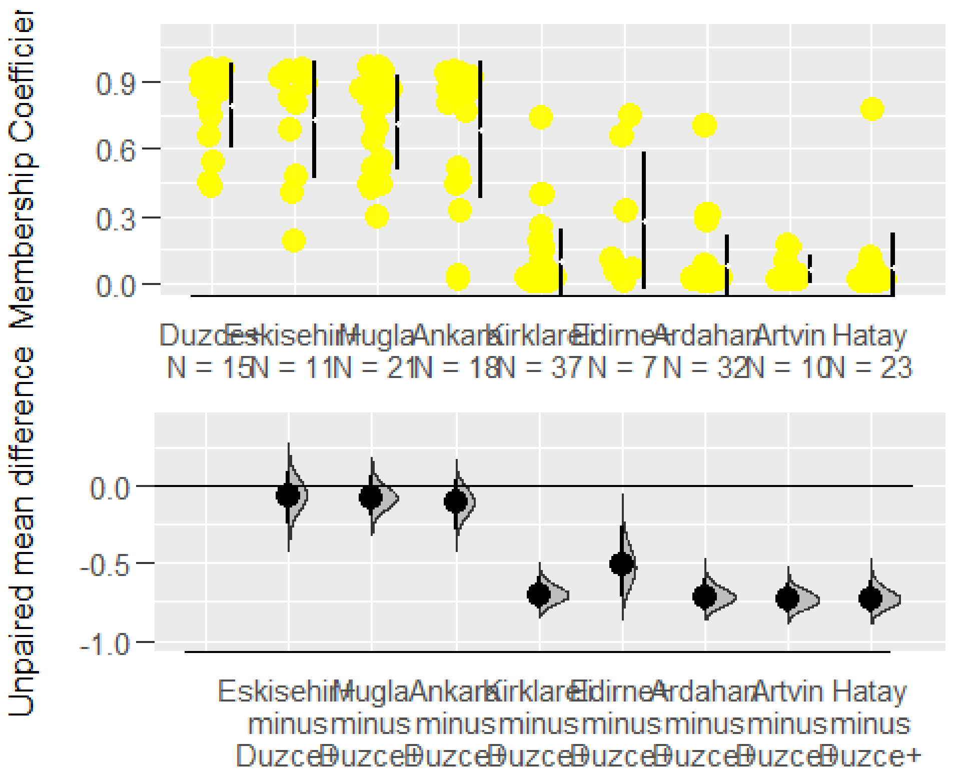


```
plot(dabest(all_provinces_iso, POPULATION, CAUCASIAN,
         idx = c("Ardahan", "Artvin", "Kirklareli", "Edirne+", "Duzce+", "Esk
isehir+", "Mugla", "Ankara", "Hatay"),
         paired = FALSE),
     theme = ggplot2::theme_gray(),
     palette = rep(c("blue"), times= 9),
     rawplot.ylim = c(0.00, 1.10),
     effsize.ylim = c(-1.00, 0.40),
     rawplot.markersize = 4,
     rawplot.groupwidth = 0.2,
     rawplot.ylabel = "Membership Coefficient",
     axes.title.fontsize = 12
)
```

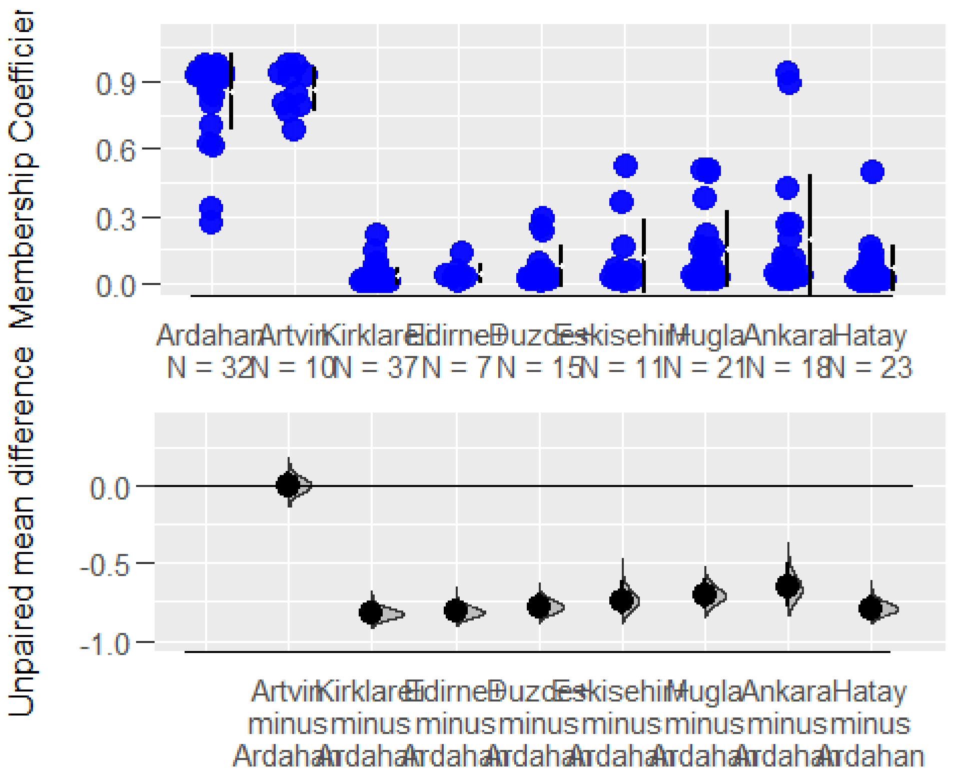


```
plot(dabest(all_provinces_iso, POPULATION, LEVANTINE,
         idx = c("Hatay", "Kirklareli", "Edirne+", "Duzce+", "Eskisehir+", "M
ugla", "Ankara", "Ardahan", "Artvin"),
         paired = FALSE),
     theme = ggplot2::theme_gray(),
     palette = rep(c("violet"), times= 9),
     rawplot.ylim = c(0.00, 1.10),
     effsize.ylim = c(-1.00, 0.40),
     rawplot.markersize = 4,
     rawplot.groupwidth = 0.2,
     rawplot.ylabel = "Membership Coefficient",
     axes.title.fontsize = 12
)
```

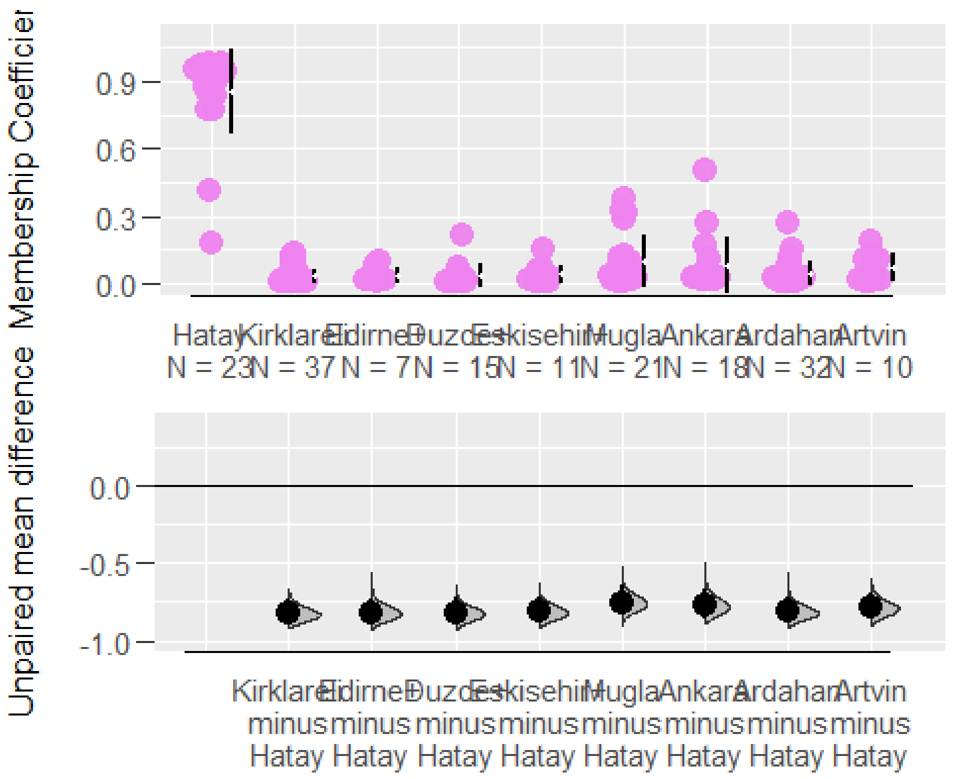


#### Supplementary

```
plot(Bitlis_mig$ANATOLIAN, pch=21, bg="gold4", cex=1.5, col="black", las=1,
  xlab="",
  ylab="Membership coefficients",
  ylim= c(0, 1.10),
  pars=list(par(mar=c(4,6,4,2))),
  main="Individual Assignments in Bitlis+")

## Warning in plot.window(...): "pars" bir grafiksel parametre değil

## Warning in plot.xy(xy, type, ...): "pars" bir grafiksel parametre değil

## Warning in axis(side = side, at = at, labels = labels, ...): "pars" bir
## grafiksel parametre değil

## Warning in axis(side = side, at = at, labels = labels, ...): "pars" bir
## grafiksel parametre değil

## Warning in box(...): "pars" bir grafiksel parametre değil

## Warning in title(...): "pars" bir grafiksel parametre değil

points(Bitlis_mig$LEVANTINE, pch=21, bg="violetred", cex=1.5, col="black")
points(Bitlis_mig$CAUCASIAN, pch=21, bg="blue", cex=1.5, col="black")
abline(h=0.5, col="black",lwd=4)
```

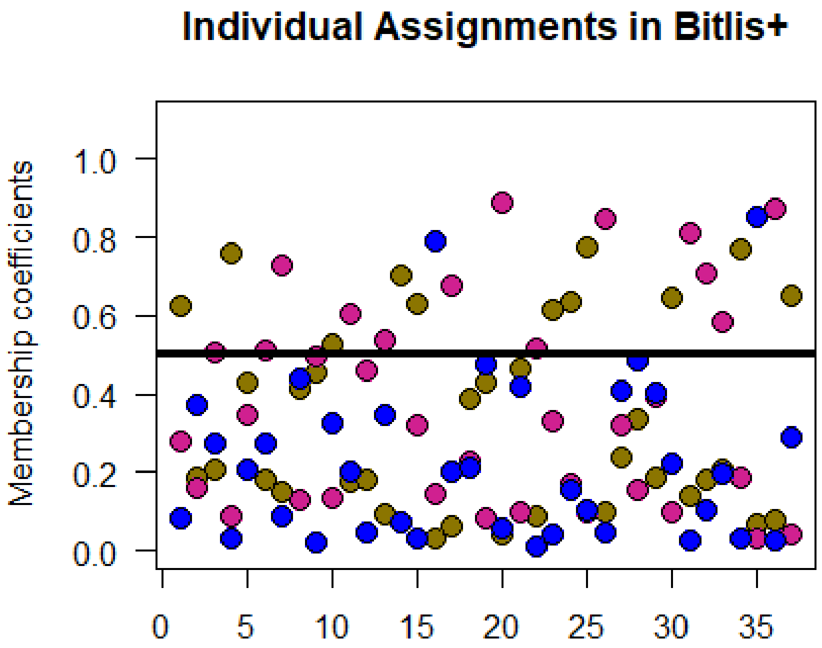


```
mean(Bitlis_mig$LEVANTINE)

## [1] 0.3690405

mean(Bitlis_mig$ANATOLIAN)

## [1] 0.347273

dunn.test(list(Bitlis_mig$ANATOLIAN, Bitlis_mig$LEVANTINE, Bitlis_mig$THRACIA
N, Bitlis_mig$CAUCASIAN),
method="bh", table=FALSE, list=TRUE, altp=TRUE)$altP.adjust

##   Kruskal-Wallis rank sum test
##
## data: x and group
## Kruskal-Wallis chi-squared = 59.0356, df = 3, p-value = 0
##
##
##                             Comparison of x by group
##                               (Benjamini-Hochberg)
##
## List of pairwise comparisons: Z statistic (adjusted p-value)
## ---------------------------
## 1 - 2 : -0.176265 (0.8601)
## 1 - 3 :  6.562506 (0.0000)*
## 2 - 3 :  6.738772 (0.0000)*
## 1 - 4 :  1.987734 (0.0562)
## 2 - 4 :  2.164000 (0.0457)*
```

```
## 3 - 4 : -4.574772 (0.0000)*
##
## alpha = 0.05
## Reject Ho if p <= alpha

## [1] 8.600852e-01 1.587321e-10 9.583830e-11 5.620928e-02 4.569651e-02
## [6] 9.534769e-06
```
